# EstroGene2.0: A multi-omic database of response to estrogens, ER-modulators, and resistance to endocrine therapies in breast cancer

**DOI:** 10.1101/2024.06.28.601163

**Authors:** Zheqi Li, Fangyuan Chen, Li Chen, Jiebin Liu, Danielle Tseng, Fazal Hadi, Soleilmane Omarjee, Kamal Kishore, Joshua Kent, Joanna Kirkpatrick, Clive D’Santos, Mandy Lawson, Jason Gertz, Matthew J. Sikora, Donald P. McDonnell, Jason S. Carroll, Kornelia Polyak, Steffi Oesterreich, Adrian V. Lee

## Abstract

Endocrine therapies targeting the estrogen receptor (ER/*ESR1*) are the cornerstone to treat ER-positive breast cancers patients, but resistance often limits their effectiveness. Understanding the molecular mechanisms is thus key to optimize the existing drugs and to develop new ER-modulators. Notable progress has been made although the fragmented way data is reported has reduced their potential impact. Here, we introduce EstroGene2.0, an expanded database of its precursor 1.0 version. EstroGene2.0 focusses on response and resistance to endocrine therapies in breast cancer models. Incorporating multi-omic profiling of 361 experiments from 212 studies across 28 cell lines, a user-friendly browser offers comprehensive data visualization and metadata mining capabilities (https://estrogeneii.web.app/). Taking advantage of the harmonized data collection, our follow-up meta-analysis revealed substantial diversity in response to different classes of ER-modulators including SERMs, SERDs, SERCA and LDD/PROTAC. Notably, endocrine resistant models exhibit a spectrum of transcriptomic alterations including a contra-directional shift in ER and interferon signaling, which is recapitulated clinically. Furthermore, dissecting multiple *ESR1*-mutant cell models revealed the different clinical relevance of genome-edited versus ectopic overexpression model engineering and identified high-confidence mutant-ER targets, such as *NPY1R.* These examples demonstrate how EstroGene2.0 helps investigate breast cancer’s response to endocrine therapies and explore resistance mechanisms.

## Introduction

Breast cancer remains the most common cause of cancer-associated death among women and is responsible for nearly 15% of all new cancer cases each year in the United States (1). This disease presents as a heterogeneous entity, with various subtypes distinguished by unique molecular profiles, guiding different treatment strategies. Among these subtypes, estrogen receptor-positive (ER+) breast cancer, characterized by the presence of estrogen receptor (ER/*ESR1*), accounts for approximately 70% of cases (2,3). Not surprisingly, targeted therapies which directly inhibit the transcriptional activity of ER, or which block the synthesis of estrogens have and will continue to be the cornerstone of interventions used to treat this disease at all stages (4–6).

The Selective Estrogen Receptor Modulators (SERMs), like tamoxifen, function as competitive inhibitors of estrogen binding to ER and also disrupt the integrity of the protein-protein interaction surfaces needed for coactivator recruitment (7). Tamoxifen is used in combination with GnRH analogues (to disrupt estrogen biosynthesis) in premenopausal women with ER-positive breast cancer (5). In postmenopausal women with this disease subtype, aromatase inhibitors (AIs), such as letrozole and anastrozole, which inhibit the peripheral conversion of androgens to estrogens are now the standard of care for frontline intervention (8). The classic models of ER-action suggest that a small molecule ligand (i.e. an estrogen) is required for the activation of the receptor. However, in cancer cells it has been shown that ER transcriptional activity can occur in a ligand independent manner secondary to the overexpression of certain coactivators (i.e. SRC3) or increased activity of signaling pathways which impinge upon the ER-coregulator complex (9). Further dysregulated expression of the expression and/or activity of coregulators such as SRC3 can also have a profound effect on the pharmacology of ER-modulators (10,11). These mechanistic insights gave rise to the idea that elimination of ER as opposed to its inhibition may be a better approach in cancer.

Emerging from these efforts was the first-generation SERD fulvestrant, whose clinical efficacy, albeit modest, validated the general approach of eliminating the receptor as a therapeutic approach (12). The first non-steroidal oral SERD, Etacstil, was developed over 25 years ago and although it demonstrated clinical efficacy its development was discontinued (13). However, the lessons learned from studying this drug informed the discovery and development of the third generation SERDs, a large number of which have been evaluated in the clinic (6). Also driving SERD development is the identification of constitutively active hotspot mutations like *ESR1* Y537S and D538G (14,15), *ESR1* fusions which exhibit neomorphic transcriptional activities and the demonstration of *ESR1* amplification in some tumors (16) (17). Elacestrant was the first SERD of this class of medicine to be FDA approved and is currently used in patients whose tumors harbor an activating ESR1 mutation (18,19). Camizestrant, giredestrant, imlunestrant and palazestrant are all in late-stage clinical trials in patients with AI resistant disease with some being evaluated in earlier settings in patients at high risk of recurrence (20–24). All of the SERDs described above have a common mechanism of action in that they induce denaturing conformational changes in ER structure which result in it being targeted to the proteasome. However, proteosome dependent degradation is also achieved by the ligand directed degrader ARV471 (vepdegestrant), a bifunctional ER ligand directed degrader (LDD) which has a receptor binding moiety and a second functionality which results in the recruitment of the cereblon E3 ligase (25). Challenging the idea that SERD activity is an absolute requirement of ER-modulators for advanced disease is the SERM lasofoxifene, has shown substantial clinical activity in patients whose tumors express *ESR1* mutations. It is likely that several oral SERDs, ARV471 and the SERM lasofoxifene will be approved clinically and thus the next challenge will be to understand how to distinguish these functionally distinct drugs and identify best clinical utility (26,27). Key to success in this regard will be to define how ER activity and pharmacology is impacted by alterations in the expression and activity of specific coregulators (coactivators and corepressors) and by the activity of PI3K(28), MAPK(29), and ERBB2 signaling pathways (30). Also important will be to define how epigenetic modifications, such as changes in DNA methylation and histone modifications, impact ER pharmacology and the response of tumors to these agents.

There are few validated *in vivo* models of ER+ breast cancer and thus cell line models have played a vital role in studies that have elucidated the molecular mechanisms that determine ER pharmacology and how pathways are dysregulated upon drug resistance. Two broad categories of endocrine resistance models have been widely utilized: consequence-mimicking and molecular-mimicking models (31). The former involves the development of cell subclones under selective pressure of chronic treatment with different endocrine therapies, *in vitro* or *in vivo*, and subsequent elucidation of the cause-and-effect relationships between specific molecular events and drug pharmacology. The tamoxifen-resistant (TamR) models and long-term estradiol deprivation (LTED) models are among the most widely used. Molecular-mimicking models, on the other hand, engineer cells with known genetic alterations identified from endocrine-resistant cases, such as *ESR1* mutations. These models, created using genome editing or ectopic overexpression, have resulted in the discovery of new biology and druggable therapeutic vulnerabilities (32). Advances in high-throughput sequencing technologies have revolutionized our ability to explore the molecular landscapes of ER modulator activity in models of endocrine therapy sensitivity and resistance. However, despite the wealth of publicly available datasets, and tools like NCBI GEO(33) and Cistrome DB(34), accessing and analyzing these data in a comprehensive manner remains challenging.

The EstroGene2.0 knowledgebase has been developed to address the limitations in processing and analyzing current datasets that probe ER-biology focusing on transcriptomic and cistromic analyses. Extending its predecessor, EstroGene1.0 (35), which focused solely on experiments analyzing treatment with estrogens. EstroGene2.0 encompasses transcriptomic and ER ChIP-seq profiling data from studies performed with a large number of different ER modulators in models of both endocrine therapy sensitive and resistant breast cancer cells. Our updated platform features an enhanced user-friendly browser, enabling researchers to swiftly access and review experimental documentation, dataset quality controls, and insights into gene regulation and ER binding associations across hundreds of curated experiments. Additionally, we have integrated an unpublished ER Rapid Immunoprecipitation Mass spectrometry of Endogenous protein (RIME) profiling data from 16 ER+ cancer cell lines, offering additional access into ER interactomes. Following the development of EstroGene2.0, we conducted a series of pilot studies to explore potential mechanisms of response and resistance to ER modulators and herein report these findings and provide examples of new testable hypothesis that can be tested experimentally.

## Materials and Methods

### EstroGene2.0 data curation

To obtain comprehensive ER modulator treatment and endocrine resistant cell model related database in breast cancer, we established a standardized curation model with three main steps. First, we conducted a literature search from the Gene Expression Omnibus (GEO database) using the combination of ER modulator compound (e.g. “tamoxifen” or “4OHT) or endocrine resistant model type (e.g. “tamoxifen resistant” or TamR) plus “breast cancer” plus the name a specific type of sequencing technology (e.g., “RNA-seq” or “RNA sequencing”) towards publications released earlier than January 2023. Secondly, we manually reviewed these articles, only literature conducting the required experiments on human breast cancer cell lines were incorporated into the EstroGene2.0 database. We curated details of 1) publications including GEO accession number, PMID, publication date and institution; 2) experimental designs including cell lines, replicates used, compound dose, duration for ER modulator treatment, resistant development procedure, model construction method and variants type (for ESR1 mutations), library preparation method and NGS sequencing platforms. All the relevant information is summarized in Supplementary Table S1.

### Website construction

The EstroGene II website utilizes Firebase (https://firebase.google.com) for its robust NoSQL database infrastructure. We developed specialized gene-based indices and stored this data structure in Firebase, which specifically provides constant time retrieval for our website. It reduces search latency, thereby enhancing the responsiveness of our platform. As a result, EstroGene II operates with higher efficiency and speed compared to its predecessor. Our frontend is crafted using Angular 16.2.0 (https://angular.io) with TypeScript, HTML, and CSS. Angular simplifies the development process with its tooling and libraries. Its efficient rendering pipelines minimize the resource footprint and enhance the speed of view updates. In addition, R Shiny was used to construct the external analysis page including mode2 and mode3 of the analysis tab. These features enable EstroGene II to handle complex interactions and maintain a reactive user experience across all user activities.

### Transcriptomic data process and analysis

For all the RNA-seq experiments, we took advantage of the recently released NCBI-generated RNA-seq count data (BETA version). Briefly, SRA runs are aligned to genome assembly GCA_000001405.15 (hg38) using HISAT2. Runs that pass a 50% alignment rate are further processed with Subread featureCounts which outputs a raw count file for each run. For Human data, the Homo sapiens Annotation Release 109.20190905 was used for gene annotation. Genes with 0 counts in each experiment were removed and DESeq2(36) was used to compute log2-fold change and adjust P values of each gene between control and targeted samples. For specific datasets lacking replicates, we generated log2-fold change of each gene by subtracting TMM normalized log2(CPM+1) values of controls from the corresponding targeted samples.

For microarray datasets, we collected the raw array files from GEO database and normalized the data with different packages according to the platform. Affy(37) and oligo(38) packages were used to process Affymetrix-based microarray data following RMA normalizations. For data generated based on Agilent platform, loess normalization was performed directly on preprocessed data were downloaded from GEO. Different version of probe ID were converted to gene ID using BioMart package. Probes representing the same gene were merged by averaging the normalized intensity. LIMMA(39) was used to compute differential expressing genes for datasets including biological replicates. For experiments without replicates, log2-fold changes were calculated by subtracting the control values from the matched samples.

For the integration of RNA-seq and microarray comparisons under the same section (e.g. short-term ER modulator treatment), the percentile of each gene within each comparison was calculated independently by ranking the E2-induced fold changes, with -100–0 as repression and 0–100 as activation and 0 as no regulation. For the merged regulatory percentile, genes appeared in below 80% comparisons of each section were filtered out. For downstream analysis, regulator prediction were performed using LISA(40). Venn diagrams were generated using jVenn(41). Data visualizations were performed using “ggpubr”(42) and “Complexheatmap”(43).

For multivariant regression modeling of short-term ER modulator treatment, comparisons defined by ER modulator treatment group of the same cell line, drug time, and time, compared to control group, we linearly scaled expression fold change of each gene (from differential gene expression analysis) to percentile rank, ranging from -100 to 100. Specifically, the most down-regulated gene is set as -100, and most up-regulated gene as 100, and genes with no change remain as 0. We selected experiments with tamoxifen or fulvestrant treatment as input for regression model. For each gene, we modeled its percentile rank by mixed effect linear regression model, using each comparison as a single observation.

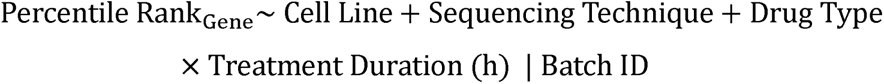

Specifically, sequencing technique is either RNA-seq or Microarray. Batch ID marks a unique experiment which consists of one to multiple experiment groups using same set of control samples. Regression model is calculated with python statsmodel package(44) (v.0.14.1). We recorded regression coefficient and significance p value for each term and interaction term for each model.

For pathway enrichment analysis, we used EnrichR(45) (https://maayanlab.cloud/Enrichr/) to calculate pathway enrichment, selecting the following datasets: KEGG (human, 2021), Reactome (2022), Gene Ontology Biological Process (2023), Gene Ontology Cellular Component (2023), and Gene Ontology Molecular Function (2023). For each dataset in ER modulator treatment, the top 5 enriched gene set, ranked by adjusted p value, were selected for presentation for each enrichment analysis. The enrichment plot was generated using gseapy(46) (v1.1.0) dotplot function, showing ratio of gene overlap with pathways as dot size, and colored by adjusted p value from pathway enrichment. For other data sets, we mainly focused on the Hallmark signature collection, and generated the enrichment bubble plots for significantly enriched pathways using adjusted p values and odds ratio.

We also used NES based method for different types of compound pathway enrichment in Fig. 3J. Briefly, we first calculated average gene percentile rankings of each gene from all comparisons treated with the specific ER modulator type. Then for each drug type, we ran gene set enrichment analysis (prerank function from gseapy v1.1.0), using MSigDB hallmark (2020) gene sets using the average gene percentile ranking as pre-ranked correlation. Other computation parameters include minimum and maximum allowed gene number from gene set and data set as 5 and 1000 respectively, permutation number as 1000, and random seed as 6. This generated a normalized enrichment score (NES) and an FDR value for each pathway and each drug type. In the heatmap, we colored drug-pathway pairs by NES values.

For clinical cohort analysis, expression data and sample metadata of POETIC trial were downloaded from GSE105777(47). Enrichment of interferon and estrogen response signatures were calculated using GSVA package(48). For *ESR1* mutant metastatic breast cancer cohorts, differentially expression genes between WT and mutant samples in each cohort was described before using padj<0.1 as the cutoff. For logistic regression modeling, GSVA enrichment scores were first calculated with up- and down-regulated genes with regulatory percentile above 60% or below -60% from 32 genome-edited and 12 ectopic overexpression models. An integrated enrichment score of each sample were calculated by subtracting the repression score from the activation score. Lme4 package(53) was used for the regression modeling using ESR1 mutation status as predictor and pROC package(54) was used to generate ROC curves.

### ChIP-seq data process and analysis

ChIP-seq raw fastq files were downloaded from GEO with corresponding SRR accession numbers. Reads were aligned to hg19 genome assembly using Bowtie 2.0(55), and peaks were called using MACS2.0 with q value below 0.05(56). For gained peaks in ESR1 mutant cells versus WT controls, we used DiffBind package(57) to intersect BED files. Intensity plots for binding peaks were visualized by Seqplots(58) using BigWig files and BED files as input. Peak visualization was conducted via WashU EpiGenome Browser(59) using BigWig files as input. For gene annotation, Binding and Expression Target Analysis(60) minus was used with 100kb as the distance from gene TSS within which peaks. Average BETA score from 16 ESR1 mutant gained peaks sets were calculated for each gene.

### Cell culture

BT474, BT483, EFM19, EFM192A, HCC1428, T47D, ZR751 & ZR7530 were cultured in in RPMI 1640 Medium (no phenol red, no L-glutamine) (Gibco # 32404014). The following supplements were added: 1mM sodium pyruvate (Gibco # 11360039) and 2mM GlutaMax (Gico # 35050038) and heat-inactivated (61)FBS was used. For BT483, human insulin (Sigma # I0516) was added at a final concentration of 0.01mg/ml. Genome-edited *ESR1* WT and mutant T47D (RRID: CVCL_0553) *ESR1* mutant cell models (62)experiments, cells were deprived in phenol-red-free IMEM with 5% CSS. CSS was purchased from Hyclone (#SH30068).

### ER RIME

Cells were seeded in 15cm plates and when 85-90% confluent, cells were double crosslinked with 2mM disuccinimidyl glutarate for 20 mins followed by 1% formaldehyde. Pulldown was performed with either ER antibody mix or Rabbit IgG as described before(61). RIME samples were digested with trypsin (Pierce) and the peptides were purified with Ultra-Micro C18 Spin Columns (Harvard Apparatus) prior to mass spectrometry analysis as previously described(62). Dried peptides were reconstituted in 0.1% formic acid for further LC–MS/MS analysis.

For LC-MS/MS Analysis, reconstituted peptides were analysed on a Dionex Ultimate 3000 system coupled with the nano-ESI Fusion Lumos (Thermo Scientific). Peptides were loaded on the Acclaim PepMap 100, 100 μm × 2 cm C18, 5 μm, 100 trapping column and separated with the EASY-Spray analytical column (75 μm × 25 cm, C18, 2 μm.) with a 5–45% acetonitrile gradient in 0.1% formic acid at 300 nL/min flow rate. The full scans were performed in the Orbitrap in the range of 400 to 1600 m/z at 120k resolution. The MS2 scans were performed in the ion trap with 2.0 Th isolation window, HCD collision energy 28% and dynamic exclusion 30 seconds.

For data analysis, Spectral .raw files from data dependent acquisition were processed with the SequestHT search engine on Thermo Scientific Proteome Discoverer 2.4 software. Data was searched using Uniprot Database Homo sapiens fasta file (taxon ID 9606 - Version June2). The node for SequestHT included the following parameters: Precursor Mass Tolerance 20 ppm, Fragment Mass Tolerance 0.5 Da, Dynamic Modifications were Oxidation of M (+15.995Da) and Deamidation of N, Q (+0.984Da). The Precursor Ion Quantifier node (Minora Feature Detector) was used for label-free quantification included a Minimum Trace Length of 5, Max. ΔRT of Isotope Pattern 0.2 minutes. The consensus workflow included peptide validator, protein filter and scorer. For calculation of Precursor ion intensities, Feature mapper was set True for RT alignment, with the mass tolerance of 10 ppm. Precursor abundance was quantified based on intensity and the level of confidence for peptide identifications was estimated using the Percolator node with a Strict FDR at q-value < 0.01.

For bioinformatics analysis, label-free quantification was used to generate peptide intensity values. The data analysis was performed using Bioconductor based R package qPLEXanalyzer. Initially, the data was normalized via within-group median scaling, where IgG served as one group and all other pulldowns constituted the other. Peptides detected in less than 25% of pulldown samples were excluded from subsequent analysis. For the remaining peptides, missing values were imputed by two methods: utilizing the minimum value in each sample (for all IgG controls and in pulldown samples for peptides found in less than half of replicates) or employing a k-nearest neighbours algorithm (for pulldown samples with peptides detected in more than half of replicates). Subsequently, a limma-based statistical analysis was conducted to identify differentially enriched proteins compared to the IgG control.

### Transient siRNA transfection

Individual *ESR1* WT and mutant T47D clones were hormone-deprived in IMEM/5% CSS for 3 days and evenly pooled for siRNA transfection. Briefly, 500,000 cells were seeded in 6-well plates in IMEM/5% CSS and forward transfected with 50 nM final concentration of ON-TARGETplus Non-targeting Control Pool siRNA (Dharmacon #D-001810-10-05), ON-TARGETplus Human NPY1R SMARTPool siRNA (Dharmacon #L-005672-00-0005), ON-TARGETplus Human RLN1 SMARTPool siRNA (Dharmacon #L-017403-00-0005), or ON-TARGETplus Human SUSD3 SMARTPool siRNA (Dharmacon #L-016811-02-0005) using Lipofectamine RNAiMAX (Invitrogen #13778) protocol for 24 or 72 hours.

### qRT-PCR

*ESR1* WT and mutant T47D cells were collected after 3 days of hormone deprivation or 72 hours post-siRNA transfection. RNA extraction was performed using RNeasy mini kit (Qiagen #74106). Reverse transcription to cDNA was performed with PrimeScript™ RT Master Mix (Takara Bio #RR036B). RT-PCR was then performed with SsoAdvanced Universal SYBR (Bio-Rad #1726275) with primers as detailed in supplemental materials (see below). The ΔΔCt method was used to analyze relative mRNA fold changes with GAPDH serving as the internal control. Statistical differences evaluated using a paired t-test. Sequences of primers are list below:

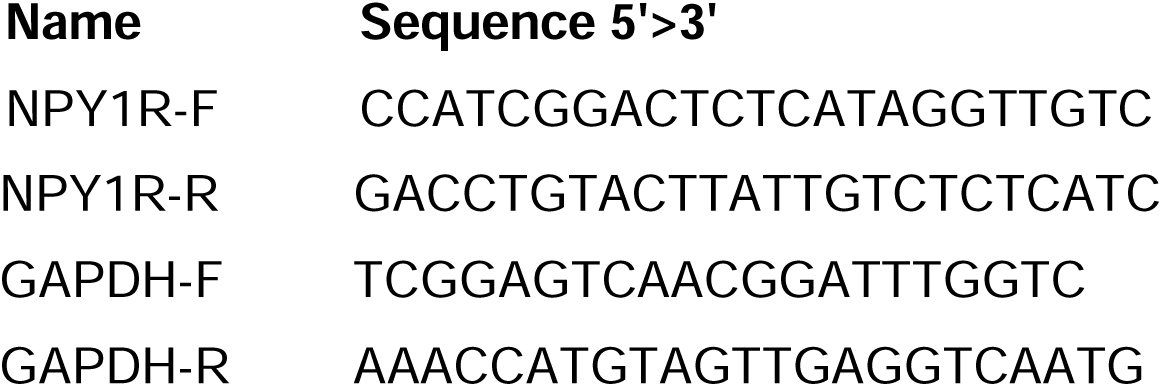

### Growth assay

Twenty-four hours after siRNA transfection, *ESR1* WT and mutant T47D cells were plated into 96-well plates at 2000 cells per well. Cell viability was measured at day 0, 2, 4, 6 and 8 and quantified with PrestoBlue HS Cell Viability Reagent (Life Technology #P50201) following the manufacturer’s protocol. Growth rate was determined as fold change compared to day 0.

## Results

### Expansion of EstroGene database to include studies of ER modulator response and resistance

To increase the clinical relevance and utility of the EstroGene database, we expanded its scope to include data generated in breast cancer cell models subjected to ER modulator treatment or designed to mimic endocrine therapy resistance. A comprehensive search was undertaken of publicly available data from the Gene Expression Omnibus (GEO) using specific keywords related to compound names (e.g., tamoxifen, fulvestrant) or resistant model types (e.g., tamoxifen resistance, *ESR1* mutation), coupled with the name of a specific technology (e.g., "RNA-seq" or "RNA sequencing"). Following this initial search, we performed manual filtration to ensure data accuracy. Our focus for endocrine-resistant models centered on tamoxifen-resistant (TamR), long-term estradiol deprivation (LTED), and *ESR1* mutation models, given their prevalence in published literature. This search strategy yielded 178 experiments from 94 studies, comprising 104 RNA-seq, 57 microarray, and 17 ER ChIP-seq experiments, supplementing the existing 92 transcriptomic and 75 ER ChIP-seq experiments from the first version of the EstroGene database. Additionally, we incorporated in-house experiments of ER RIME (Rapid Immunoprecipitation Mass Spectrometry of Endogenous proteins )(65) proteomic profiling across 16 ER+ cancer cell lines (15 breast cancer and one endometrial cancer) representing an unbiased survey of ER-associated protein complexes using IP-mass spectrometry. The EstroGene2.0 database now encompasses multi-omic profiling data from 361 experiments across 212 studies, spanning 28 cell lines and publications from 2004 to 2023 (Fig. 1A and Supplementary Table S1).

**Figure 1.**
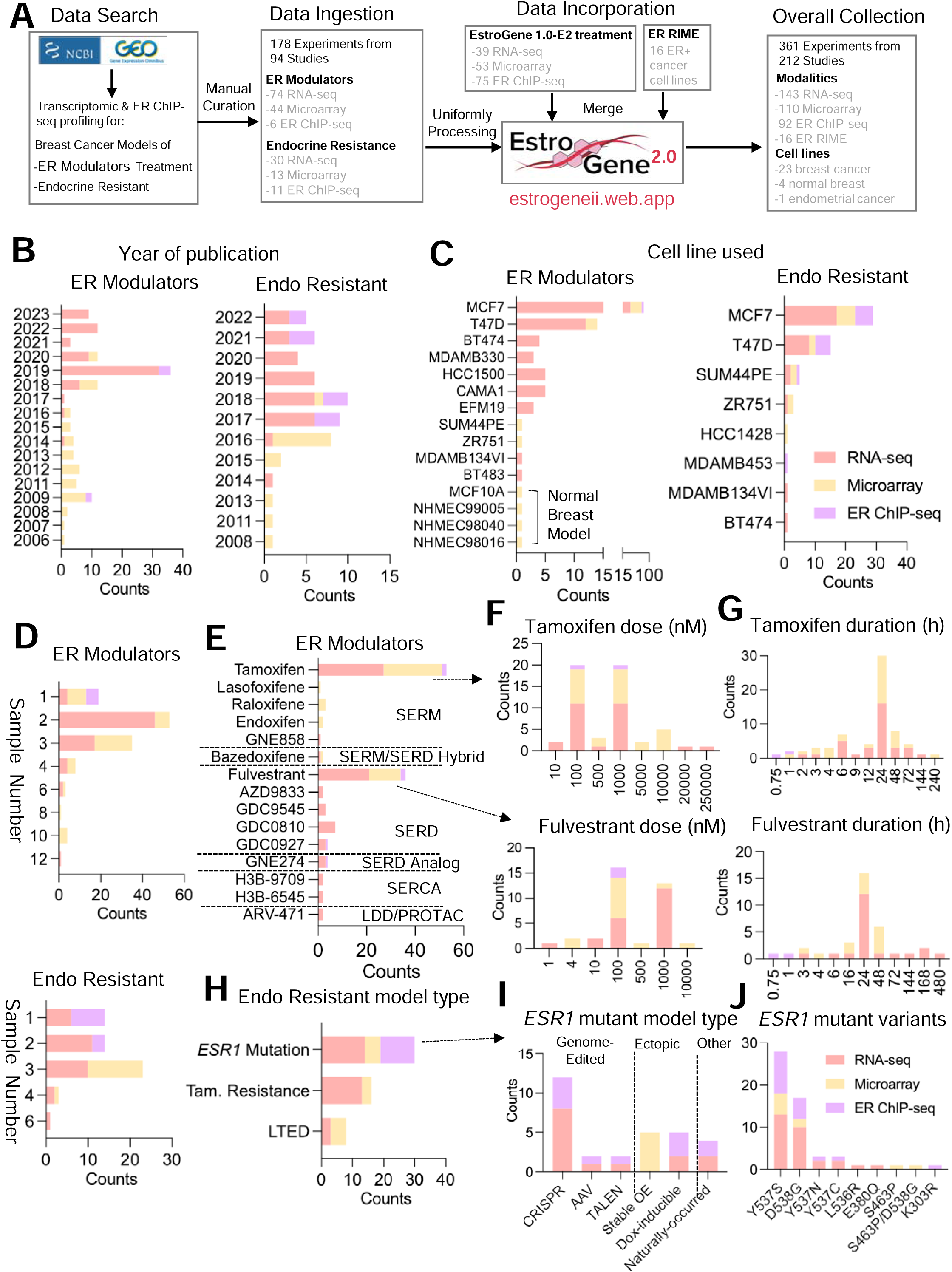
Expansion of EstroGene database towards ER modulator response and resistance profiles. A. A flow chat depicting the process for the EstroGene2.0 database collection. B-J. Stacked histogram showing the metadata separated by technologies and two major model types, ER modulator treatment and endocrine resistance (Endo Resistant), across all the curated datasets related to year of dataset publication (B), cell line used (C), replicates used (D). Within ER modulator treatment collection, further separation by compound types (E), dose (F) and duration (G) for tamoxifen and fulvestrant are shown. Within endocrine resistant models, further subgroup by specific model types (H), ESR1 mutant cell model types (I) and specific variants (J) are shown.

Similar to observations from EstroGene1.0 (35), publicly available data from studies performed after 2017 was for the most part generated using next-generation sequencing technologies, while microarray was more prevalent in earlier studies (Fig. 1B). Notably, most studies (78.6% RNA profiling and 77.4% ChIP seq) were performed in the ER-positive MCF7 and T47D cell lines (or endocrine therapy resistant models thereof), and while this ensured that the data generated were robust it is also expected to introduce unavoidable biases (Fig. 1C). Most of the transcriptomic profiling experiments employed biological duplicates (39.8%) or triplicates (36%), whereas 82.3% of ER ChIP-seq experiments included only one sample (Fig. 1D).

For experiments using ER -modulators, we ingested data from six categories of compounds (SERM, SERD, SERM/SERD Hybrid, SERCA, LDD/PROTAC and SERD Analog), with tamoxifen (42.7%) and fulvestrant (29%) being the most frequently used SERM and SERD, respectively (Fig. 1E). It is noteworthy that the majority of studies used doses of tamoxifen or fulvestrant ranging from 100 nmol/L to 1000 nmol/L (Fig. 1F), considerably higher than the reported physiological serum peak concentrations from previous pharmacokinetic studies in animal models and in humans (66,67) (68). This raises the concern that some of the changes noted may be related to off-target activities of these drugs. Transcriptomic profiling experiments typically involved longer compound exposure, with a median treatment duration of 24 hours, while all ER ChIP-seq experiments employed a maximum treatment duration of only 1 hour (Fig. 1G).

We gathered 30, 16, and 8 experiments from *ESR1* mutation, TamR, and LTED endocrine-resistant models, respectively (Fig. 1H). TamR models were profiled as early as 2008, whereas studies on *ESR1* mutation models have increased since 2013 (Supplementary Fig. S1A). Researchers have employed various strategies to model hotspot *ESR1* mutations in breast cancer cells, including genome editing approaches such as CRISPR/Cas9 or recombinant adeno-associated viral system (rAAV, 53.3%), and conventional ectopic overexpression (33.3%). Only one study revealed cell lines with natural occurrence of *ESR1* mutations under long-term estradiol deprivation. In genetically engineered cells, the *ESR1* Y537S and D538G hotspot variants were the most frequently modeled, reflecting their high frequency detected in patient samples (49) (Fig. 1J). The impact of clonal selection of engineered cells and how this introduces biases into the resultant transcriptomes is largely unanswered in studies performed to date. Regarding TamR models, the development procedures vary, with most models using *in vitro* selection with 100-1000 nmol/L 4OHT for over six months, while one study employed in vivo selection followed by harvesting of tumor cells for downstream characterization (Supplementary Fig. S1B-S1C) (69).

### The EstroGene2.0 browser

With the curated datasets, we downloaded and performed unified data processing (detailed in the Methods section). We amalgamated these datasets into a singular web server platform, facilitating tailored data exploration and visualization for researchers, particularly those without specialized bioinformatics skills. The EstroGene2.0 web server categorizes datasets into five distinct biological sections: treatment with an estrogen (from EstroGene1.0), ER-modulator treatment, TamR, LTED, and ESR1 mutation models. Building upon the foundation of the original EstroGene browser, this updated version introduces significant feature enhancements detailed below.

The web server is structured around two primary functional modules: ‘metadata’ and ‘analysis’ (Fig. 2). The metadata section offers detailed insights into individual experiments, including sample-level details and experimental quality controls. Users can navigate through specific sections and refine their search parameters by employing filters such as modalities, cell lines, compounds (ER-modulator), or mutation variants (*ESR1* mutation) (Supplementary Fig. S2A). Upon identifying experiments of interest, users can delve deeper into sample-level documentation by accessing corresponding experiment IDs. For transcriptomic profiling, a searchable framework is provided alongside the experimental details. Users can retrieve raw normalized gene expression values from the "Expression Matrix" tab by inputting a gene of interest (Supplementary Fig. S2B), while the "Differential Expression" tab furnishes log2 fold change values with adjusted p-values for all reasoned comparisons within a specific experiment (Supplementary Fig. S2C). This functionality enables users to assess the suitability of chosen experiments or models for addressing their research queries. Additionally, principal component analysis is available for each experiment, aiding in the evaluation of variations across replicates and biological intervention (Supplementary Fig. S2D). The complete sample-level normalized expression matrix and differential expression output are accessible for download via the "Download" tab (Supplementary Fig. S2E). For ER ChIP-seq datasets, users can access ER peak numbers and mapped reads for each sample (Supplementary Fig. S2F). Notably, a sample-level genomic track view feature allows direct visualization of ChIP signal intensity and the distribution of the user’s locus of interest through integration with the WashU Epigenome Browser (Supplementary Fig. S2G). Additionally, sample-level peak calling files and normalized read counts are available for download (Supplementary Fig. S2H).

**Figure 2.**
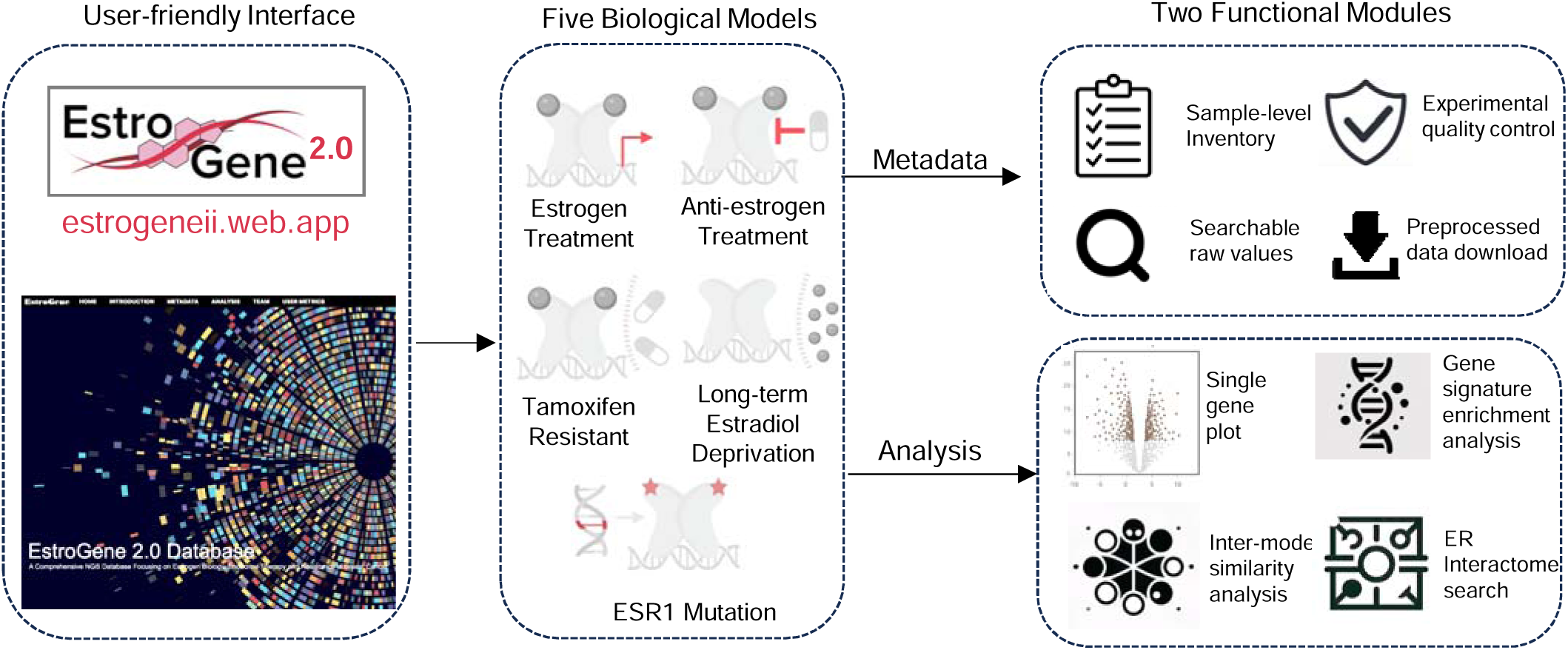
The EstroGene2.0 browser. A flow chart summarizing the conception of EstroGene2.0 website construction, including interface design, five biological models and specific functions for metadata and analysis modules.

The analysis module encompasses four modes (Supplementary Fig. S3A). The single gene visualization mode (Mode1) continues the same strategy used in the Estrogene1.0 browser, whereby users can input a gene of interest (GOI) and generate volcano plots to visualize concordance of gene expressional changes associated with the specific conditions across all the pre-computed comparisons (E2: n=149; ER modulators: n=112; TamR: n=15; LTED: n=16; ESR1 mutation: n=46, Supplementary Fig. S3B). A filter function allows users to tailor analyses towards specific experimental contexts. Users can access the original GEO deposition page by clicking on a data point in the plot. In Estrogene2.0, we additionally provide a percentile distribution plot as an alternative approach, which projects the gene regulatory percentile in a -100 to +100 window (100 represents the value of the topmost repression or activation normalized to all the processed genes in each comparison) within each comparison and merges all the results together, which offers further quantitative estimation of regulatory trend and consistency across experiments. As shown in Supplementary Fig. 3C, a well studied ER inducible gene *GREB1* showed predominant enrichment in the range of +80 to +100 with a few exceptions below 0 in *ESR1* mutant cells, depicting the quality of this visualization method alongside a representation of noted experimental variations. For the estrogen treatment and *ESR1* mutation sections, the analysis pages include targeted ER genomic binding plots based upon ER ChIP-seq data. The TSS (Transcriptional Start Site) region view summarizes peak intensities within a -/+ 200 kb range of the input gene’s transcriptional start site (Supplementary Fig. S3D), while the genomic track view redirects users to the WashU Epigenome Browser for more detailed visualization (Supplementary Fig. S3E). While the dot represents the center of each ER peak in the TSS proximal region view, users can directly assess the full genomic coordinates by hovering the mouse over a dot, facilitating the selection of regions of interest for downstream applications such as ChIP-qPCR. The gene signature enrichment analysis mode (Mode2) enables an estimation of enrichment changes of a user-defined multiple-gene signature across all the comparisons in a volcano plot format. The backend calculates geometric mean of all the customized input genes, estimates the fold change and associated adjusted p value of each comparison and separately plots by estrogen treatment, ER-modulator treatment and endocrine resistance models with specific condition filters. As shown in Supplementary Fig. S3F, inputting our previously generated EstroGene estrogen response signature (35) resulted in nearly all positive values in all the E2-treatment experiments, confirming the robustness of this signature. In Mode 3, the cross-model pattern and similarity analysis provides a holistic examination of a user’s input gene. Firstly, it showcases the regulatory trend and consistency across four biological models in a side-by-side comparison. Secondly, it identifies other genes displaying similar regulation patterns within each section, scored by Pearson correlation of regulation percentile. Thirdly, it determines the intersection of significantly similar genes across different models. For instance, using *GREB1* as an illustration, we observe a bimodal distribution of its regulatory percentile in E2 treatment and *ESR1*-mutant models (close to +100%) and ER modulator treatment experiments (close to -100%) and *vice versa* for an ER repressed gene *CCNG2* (Supplementary Fig. S3G). Notably, *ESR1*-mutant cells exhibit a considerably higher number of genes with similar regulatory trends as *GREB1* (Supplementary Fig. S3H). Finally, in Mode 4, we integrate in-house RIME experiments from 16 ER+ cancer cell lines cultured in full medium. Users can ascertain whether a target of interest is an ER interaction partner and determine the most suitable cell line for studying this interaction. FOXA1, an ER interactive pioneer factor, is presented as a default entry (Supplementary Fig. 3I).

In summary, the EstroGene2.0 browser not only broadens the scope of biological contexts and experimental modalities but also significantly enhances analytical functionality and diversity. This update provides an accessible, rapid, and comprehensive overview of cis- and trans regulation in estrogen/ER modulator response and resistance in breast cancer.

### Diversity of short-term transcriptomic response to ER-modulators

The EstroGene2.0 database not only represents a user-friendly accessible webserver for customized investigation, but also produces a harmonized large data set. We utilized this opportunity to address several crucial questions in the field, particularly focusing on data congruence. We first analyzed the impact of short-term treatment with ER modulators, based upon 123 pre-computed comparisons across 51 individual studies. Within each comparison, genes were ranked based on fold change normalized to corresponding controls, with regulatory percentiles scaled from -100 to +100, representing the spectrum from topmost down-regulation to up-regulation. Given the prevalence of tamoxifen (n=63 comparisons) and fulvestrant (n=29 comparisons) experiments in our dataset and the long-standing history in breast cancer therapeutics of both drugs, we initiated our analysis by directly comparing their effects. We observed a significantly higher variation in overall gene regulation in tamoxifen-treated experiments compared to fulvestrant-treated (Fig. 3A), indicating a more heterogeneous response to tamoxifen. To elucidate the factors driving this variation, we constructed a multivariate regression model incorporating drug type, dose, duration, sequencing modality, and cell line for each gene across all tamoxifen and fulvestrant experiments. In addition to drug types, dose emerged as one of the most influential predictors for post drug treatment transcriptomic alterations (Fig. 3B). Previous studies have highlighted the dose-dependent nature of interventions with ER modulators (66,67). Therefore, we initially assessed the regulatory percentile correlation between experiments using 1 μmol/L and 100 nmol/L doses of tamoxifen and fulvestrant. Fulvestrant exhibited a robust positive correlation (R=0.78) between the two doses (Supplementary Fig. S4A), whereas tamoxifen displayed a weaker correlation (R=0.21) (Supplementary Fig. S4B), indicating a notable dose-related discrepancy. Further stratification identified 193 and 169 genes that were more susceptible to repression by 1 μmol/L and 100 nmol/L doses of tamoxifen (Supplementary Table S2), respectively, including notable candidates such as *TFF3* and *IRX4* (Supplementary Fig. S4C). Pathway enrichment analysis revealed that 100 nmol/L tamoxifen could more effectively block estrogen response signaling, while 1 μmol/L tamoxifen preferentially suppressed TNFα/NFkB, hypoxia, and TGF-β pathways (Supplementary Fig. S4D). These analyses underscore the importance of dose in considering the pharmacology of ER-modulators and explain in part the large intrinsic variation among data sets describing the activities of these drugs in different models.

**Figure 3.**
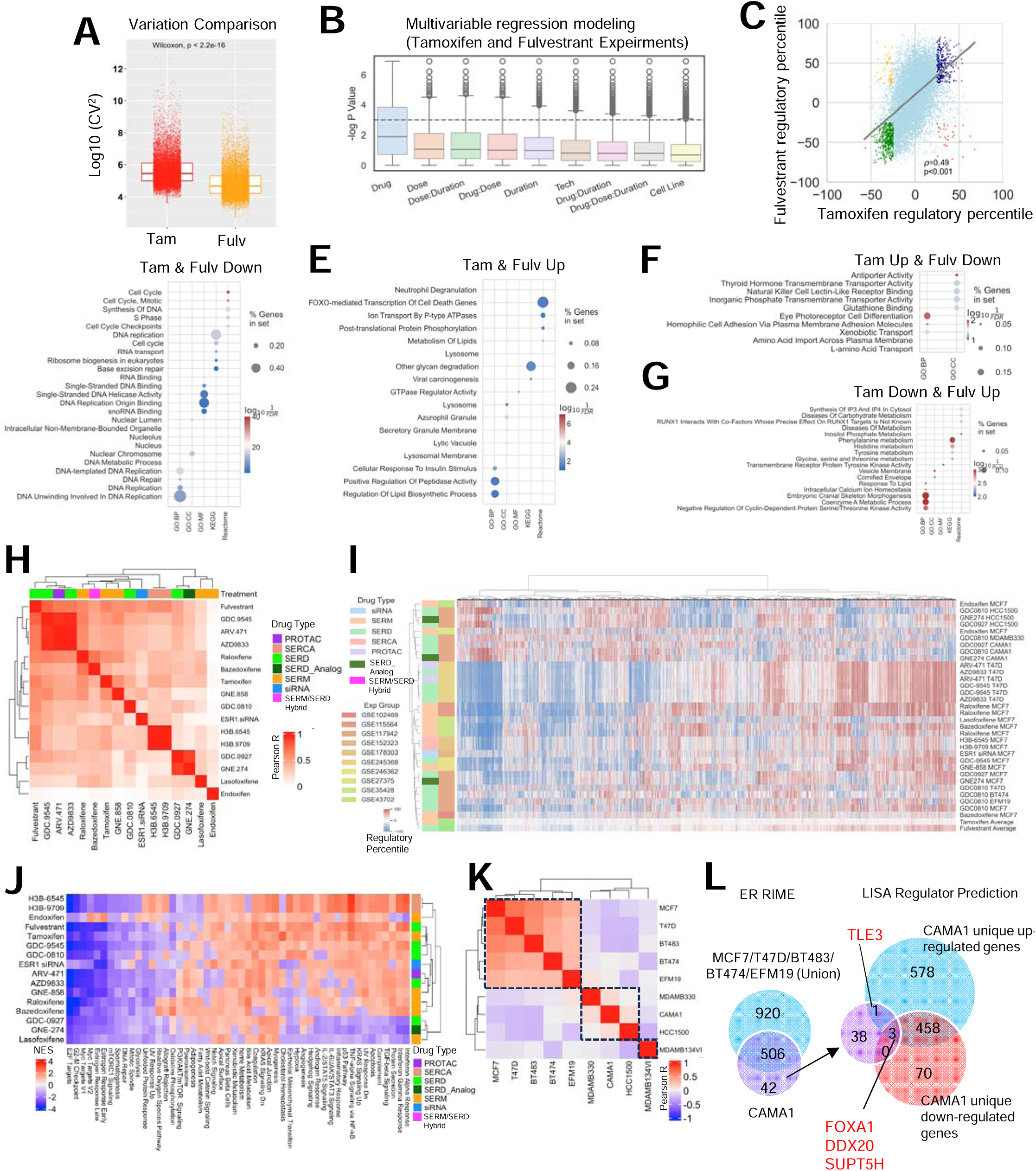
Diversity of short-term ER modulator transcriptomic response. A. Boxplots depicting the log10 (coefficient variations)^2 as a measurement of data set variation within all the tamoxifen and fulvestrant comparisons. Mann Whitney U test was used. B. Distribution of log p values of each variable from mixed effect multivariant linear regression model of gene regulatory percentiles tile from -100% to 100% with short-term tamoxifen or fulvestrant treatment experiments, among all genes. C. Scatter plot showing Pearson correlation of average regulatory percentiles of each gene from tamoxifen and fulvestrant treatment experiments. Highlighted dots include consistently regulated genes with average fold change percentile ≥25 and ≤-25 in both tamoxifen or fulvestrant treatment experiments (‘Tam-up & Fulv-up’, blue and ‘Tam-down & Fulv-down’, green) as well as inconsistently regulated genes with average regulatory percentile ≥25 and ≤-25 in one of the compounds (‘Tam-up & Fulv-down’, red and ‘Tam-down & Fulv-up’, yellow). D-G. Dot plots showing enriched pathways in genes which were ‘Tam-up & Fulv-up’ (D), ‘Tam-down & Fulv-down’ (E), ‘Tam-up & Fulv-down’ (F), and ‘Tam-down & Fulv-up’ (G), using five different databases. H and K. Heatmaps illustrating the Pearson correlation coefficients of global gene regulatory percentile between each of the two compounds (H) and cell lines (K) with unsupervised clustering. I. A heatmap showing the regulatory percentile of 11,191 genes in each of the indicated experiments and average values of fulvestrant and tamoxifen. Cell lines and compound names are labeled with color coded experiments and drug types. J. A heatmap showing GSVA normalized enrichment score (NES) of average drug-caused enrichment changes from 50 MSigDB Hallmark signatures with unsupervised clustering. L. Left: Venn diagram showing the overlapping of ER interactors from RIME experiment (log2FC>5, padj<0.05 to IgG) between CAMA1 and union of MCF7, T47D, BT483, BT474 and EFM19. Right: Overlapping of CAMA1 unique ER interactors and predicted regulators from CAMA1-specific up- and down-reglated genes. Consistent targets are labelled.

We next integrated the regulatory effects of tamoxifen and fulvestrant. Analyzing the gene-level correlation of average regulatory percentiles between tamoxifen and fulvestrant treatments revealed a positive association (R=0.49). Notably, tamoxifen exhibited a narrower average distribution range, confirming the higher intrinsic variation (Fig. 3C). To delve deeper, we examined genes showing concordant or discordant changes between these two compounds. Pathways associated with cell cycle and DNA replication were downregulated in both models (Fig. 3D), aligning with the anti-proliferative activities that are attributable to SERMs and SERDs. Among commonly upregulated genes, enrichment was observed in FOXO-mediated cell death, glycan degradation, and lysosome pathways, indicating an acute elevation in adaptation to external stress stimuli (Fig. 3E). Notably, genes specifically upregulated with tamoxifen but downregulated with fulvestrant were enriched in transport activity (Fig. 3F), while genes upregulated with fulvestrant but downregulated with tamoxifen were involved in amino acid metabolism (Fig. 3G). Some of these differences may relate to the partial agonist activity of tamoxifen manifest in some cell/promoter contexts but may also be a manifestation of the difference in inhibiting ER action (SERM) vs eliminating ER expression (SERDs).

Tamoxifen and fulvestrant are currently used for the treatment of ER-positive breast cancers. However, of late there have emerged several new classes of ER-modulators among these are (a) oral SERDs (camizestrant (AZD9833), giredestrant (GDC9545), imlunestrant, and palazestrant (b) SERMs (lasofoxifene) (c) SERM/SERD hybrids (i.e. elacestrant and bazedoxifene), (d) Ligand directed degraders (LDDs) (ARV471), (e) ligands which inhibit ER through covalent binding to the receptor (SERCAs) and (f) ER non-degrader structurally similar to SERD (SERD analog) (6). This prompts the question as to the mechanisms by which these drugs distinguish themselves at the level of transcriptional activity and how and if this could impact their clinical use. In our analysis, we evaluated four types of compounds (SERMs, SERDs, a SERCA, and ARV-471) from 10 individual studies, using *ESR1* siRNA knockdown as a reference. Pairwise average gene regulatory percentile correlations of each compound revealed an overall positive correlation, with R values ranging between 0.2 and 0.7, except for the SERM endoxifen, which showed no correlation with other compounds (Fig. 3H). ARV-471 and a number of SERDs (including fulvestrant, GDC9545, and AZD9833) were generally more similar to each other, whereas SERMs/SERM SERD hybrids (including tamoxifen, raloxifene, and bazedoxifene) were distinctly clustered from each other. Additionally, SERCA H3B-6545 and H3B-9707 showed a more similar regulatory landscape. Unsupervised clustering of all genes recapitulated an overall similar regulatory trend, despite data from the same studies tending to be more alike (Fig. 3I). To investigate unique pathway regulation by each drug type, we calculated normalized enrichment scores (NES) using the MSigDB Hallmark database (Fig. 3J). Notably, we observed consistent inhibitory effects of all drugs towards estrogen response and cell cycle-related signatures, as expected. The two SERCAs (H3B-6545, H3B-9709), endoxifen and lasofoxifene, specifically inhibited metabolic pathways involved in peroxisome and fatty acid metabolism. ARV-471 and AZD9833 exhibited additional blockade towards notch signaling, Wnt-β catenin, and apical surface cell adhesion function, which were not discerned with fulvestrant. Furthermore, a strong repression of interferon response signaling was observed in with second generation SERDs GDC0927, GNE274, and with lasofoxifene. Overall, this analysis highlights the functional diversity exhibited by different types of ER-modulators despite their similar antiproliferative activities in cellular models.

Context-dependent effects of ER modulators was observed by gene level clustering in a few cell lines (Fig. 3I), namely MDA-MB-330, HCC1500, and CAMA1. These cell lines displayed distinct gene regulatory patterns, with some genes even showing opposite regulation (Fig. 3I). This effect was further reflected in a cell-line based clustering, where a major cluster (MCF7, T47D, BT483, BT474, and EFM19) and a minor cluster consisting of MDA-MB-330, CAMA1, and HCC1500 were segregated (Fig. 3K). Additionally, MDA-MB-134VI, an invasive lobular carcinoma cell line, exhibited a distinct response pattern distinguished from other models, as we reported previously (70). It is likely that these differences in response to ER-modulators reflects the impact of these ligands on receptor structure and how this regulates the interaction of the receptor with functionally distinct cell-line specific coregulators. To explore this possibility, we conducted ER RIME profiling in 16 ER+ cell lines, including all five in the major cluster and CAMA1 in the minor cluster. Among the 15 breast cancer lines, we identified 95 proteins that consistently interact with ER including the canonical coregulator NCOA5 (Supplementary Fig. S4E and Supplementary Table S3). Further intersection analysis identified 42 CAMA1-specific ER interactors (Fig. 3L, Supplementary Fig. S4F and Supplementary Table S4). We inferred the upstream regulators controlling genes uniquely up and down-regulated in CAMA1 normalized to the major cluster cell lines using LISA (40) (Supplementary Table S4) and overlaid the unique ER interactors with the predicted regulators in CAMA1 (Fig. 3L and Supplementary Fig. 4G-4H). We identified four overlapping targets, including FOXA1, DDX20, and SUPT5H in both groups, and TLE3 as a unique ER interactor associated with CAMA1-specific up-regulated genes in response to ER modulator treatment. Of note, TLE3 has previously been characterized as an ER co-repressor (71). This suggests that ER repression in CAMA1 cells might rely heavily on TLE3, and dictate a subset of transcriptional responses in this cell line.

### Transcriptomic heterogeneity of endocrine resistant breast cancer cell models

The substantial diversity in response to ER-modulators underscores the heterogeneous mechanisms that impact receptor pharmacology and which underly resistance. Consequently, we classified the transcriptomic features of models with resistance to different types of ER-modulators. Our primary focus was on models of Tamoxifen Resistance (TamR), Long-Term Estrogen Deprivation (LTED), and those which harbored *ESR1* mutations, where the number of datasets collected provided sufficient statistical power to draw solid conclusions (n=15 comparisons for TamR, n=16 for LTED, and n=46 for ESR1 mutation). Correlation analysis utilizing average regulatory percentages from all comparisons of these three models, along with E2 regulation as a control, revealed distinct shifts in estrogenic signaling amongst the three models (Fig. 4A): 1) *ESR1*-mutant cells exhibited a high similarity to E2 treatment (R=0.46), indicative of their well-characterized ligand-independent ER activation. 2) LTED displayed a strong negative association with E2 stimulation and *ESR1* mutant models, underscoring the selection of clones with a repression of ER signaling, potentially making their growth less dependent on ER. 3) Transcriptomic features of TamR models showed minimal association with global estrogen response. Examining the overlap of genes enriched in the top 10% percentile of up- or down-regulated targets across at least 40% of comparisons for each model highlighted the distinct nature of different endocrine-resistant types (Fig. 4B and Supplementary Table S2) with minimal to no overlap of regulated genes. *LIN7A* emerged as the only consistent downregulated target, while ER target genes *GREB1*, *PGR*, and *CA12* were shared down-regulated targets between TamR and LTED models. Unsupervised clustering of Hallmark signature alterations within each model normalized to their corresponding controls (i.e., parental cells for LTED and TamR, isogenic *ESR1* wild-type models for ESR1 mutants) further revealed divergent functional shifts that correlated with the resistance phenotype (Fig. 4B). On average, *ESR1* mutant cells strongly upregulated the expression of genes associated with estrogen signaling, cell cycle progression E2F and G2M checkpoints, as well as Wnt-β-catenin signaling. In contrast, LTED and TamR models showed a decline in estrogenic and cell cycle progression signature enrichment but gained unique features. LTED models exclusively exhibited enhanced metabolic functions such as fatty acid metabolism, bile acid metabolism, and peroxisome activities, while TamR lines displayed a more diverse range of gain-of-functions with several exclusively enriched in proinflammatory signatures such as TNFα signaling via NFκB, IL6-JAK-STAT3, and inflammatory responses (Fig. 4C). Notably, the evident repression of E2 response signatures but lack of a global E2 regulation association in TamR models suggests that only a subset of canonical estrogen response program was impacted, which might be sufficient to trigger or maintain tamoxifen resistance.

**Figure 4.**
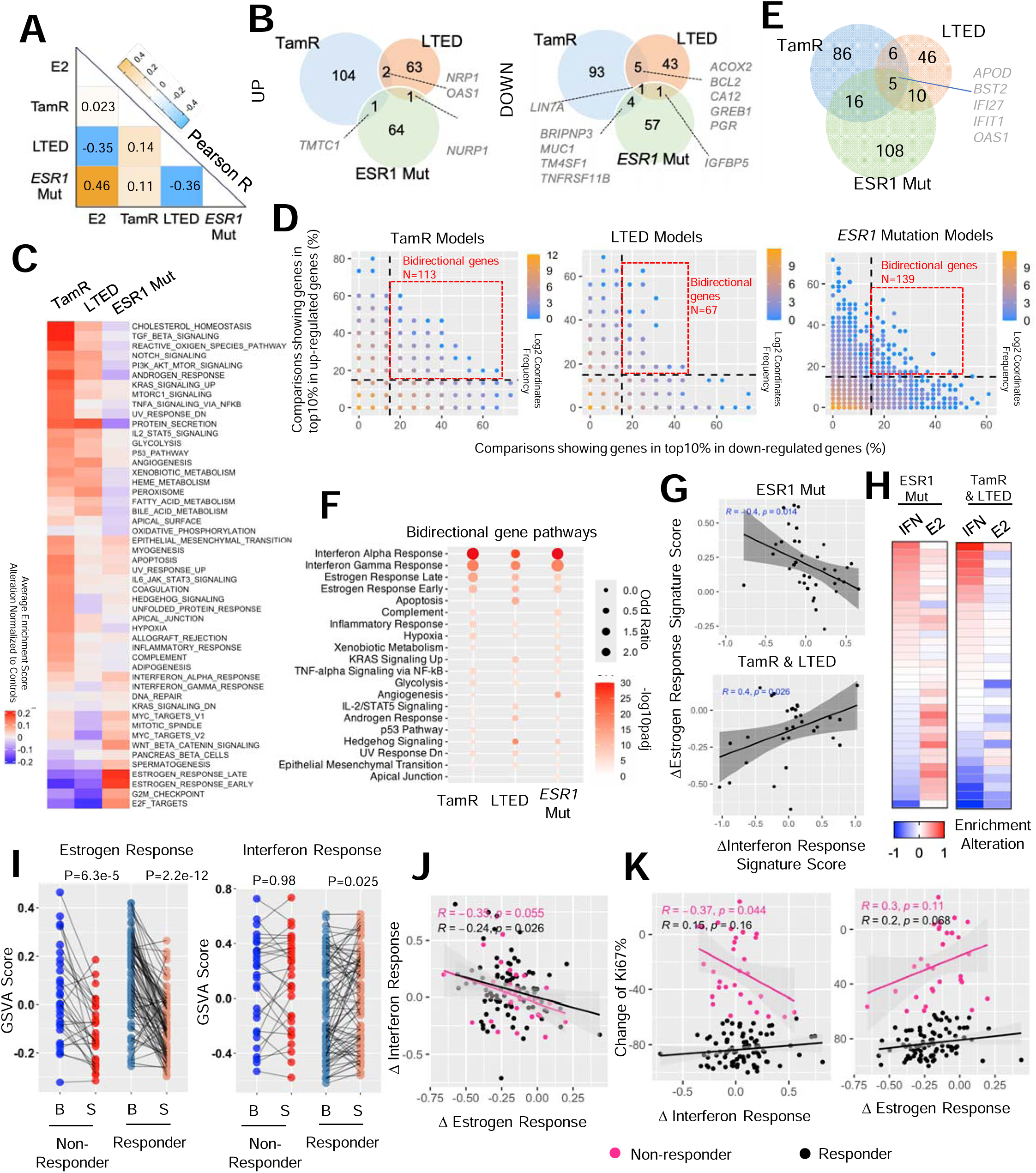
Transcriptomic heterogeneity of endocrine resistant breast cancer cell models. A. A heatmap representing the Pearson correlation coefficient between average regulatory percentile from each of the two endocrine resistant models and estrogen treatment experiments. B. Venn diagrams showing the overlay of genes that fall into top 10% up- and down-regulation and consistent in at least 40% comparisons of each endocrine resistant models. C. A heatmap depicting the average alteration of GSVA enrichment scores of each cell models normalized to the corresponding controls from 50 MSigDB Hallmark signatures with unsupervised clustering. D. Scatter plots showing the correlation of percentage of each gene that falls into top 10% up- and down-regulated parts. Color code indicates the density of each proportion. Bi-direction genes are highlighted in red frame and defined as genes showing top10% activation and repression simultaneously in at least 15% comparisons. E. A Venn diagram depicting the overlapping of bidirectional genes from the three endocrine resistant models in D. F. Dot plot showing the significantly enriched Hallmark pathways from bidirectional genes of each mode. Pathways are prioritized towards commonly changed ones. G. Scatter plot showing the Pearson correlation between average shift of interferon response signatures (mean of interferon response α and γ) and estrogen response signatures (mean of estrogen response early and late) in ESR1 mutant models and TamR/LTED models respectively. H. Heatmaps showing the average shift of interferon response signatures and estrogen response signatures of each comparison indicated in G as rows. I. Line plots depicting the change of interferon response signatures (right) and estrogen response signatures (left) between baseline and surgical samples in responder and non-responder from POETIOC clinical trial. Wilcoxon signed rank paired sample test was used. J and K. Scatter plots representing the Pearson correlation between treatment-caused changes of interferon response signatures and estrogen response signatures (J) as well as each of them towards treatment-induced changes of KI67% (K) of each patient separated by responders and non-responders.

Correlating the probability of a gene being in the top 10% of up- and down-regulated genes revealed a non-linear negative association, indicating that most genes are regulated in a single direction. The majority of these genes showed a correlation below 20%, underscoring significant variation among these experimental models (Fig. 4D). Intriguingly, we identified a subset of bidirectionally regulated genes in each model within the top 10% of both upregulated and downregulated targets and in at least 15% of comparisons (Fig. 4D and Supplementary Table S2). Five genes were shared among all three models, with four of them being involved in innate immune response (*BST2*, *IFI27*, *IFIT1*, and *OAS1*) (Fig. 4E and Supplementary Fig. S5A). Furthermore, unbiased pathway analysis consistently revealed enrichment of interferon α and γ response signatures in these bidirectional gene sets from all three models, and this divergence was not associated with the cell line background (Supplementary Fig. S5B). Given that the shift in estrogen response is a prominent feature of these endocrine-therapy resistant models, we tested whether the noted heterogeneity in the innate immune response signatures and estrogen response are linked. We separately analyzed this correlation in *ESR1*-mutant and TamR/LTED models, given the previously characterized nature of activated or repressed estrogen response programs, respectively. Surprisingly, we found that the degree of change in estrogen response signature showed a negative association with an interferon response signature in both models (Fig. 4G), where interferon signatures were only elevated in *ESR1*-mutant models with weak estrogen activation and TamR/LTED models with weak estrogen repression (Fig. 4H). These results highlight the possibility that the innate immune cascade serves as a universal negative mediator of estrogen signaling in endocrine-therapy resistant breast cancer.

We examined the relationship between response to endocrine therapy and innate immunity using data from the recently published POETIC phase III neoadjuvant endocrine therapy trial in postmenopausal women (n=115)(72). We compared baseline and surgical samples after treatment with an aromatase inhibitor revealing a significant overall decrease in estrogen response signatures in both responders (n=85) and non-responders (n=30), albeit with considerable inter-patient heterogeneity (Fig. 4I). Intriguingly, interferon response signatures were increased within the responder group (Fig. 4I). Moreover, we found that the degree of estrogen response and interferon response associated with endocrine therapy were inversely correlated, regardless of patient outcome (Fig. 4J), suggesting either mutual or mono-directional inhibitory effects between the two signals and consistent with the observation in cell models. Importantly, we observed a numerically significant inverse correlation between interferon response and changes in tumor Ki67 levels associated with treatment, specifically in non-responders (Fig. 4K). In conclusion, these analyses unveiled that endocrine therapy may trigger interferon response signatures which is associated with greater endocrine response. This sheds light on potential therapeutic avenues for overcoming endocrine resistance in breast cancer patients.

### Concordance among *ESR1*-mutant breast cancer cell models

Among all the endocrine resistance model types, hotspot *ESR1-*mutant models have the largest data set available. These mutations have undergone extensive characterization over the past decade, facilitated by the establishment of a variety of different cell models (64,73–75). However, the majority of published studies have drawn their conclusions from a single cell model, with only a few conducting cross-validation experiments through collaborative efforts (49,52). Leveraging our comprehensive collection of datasets we aimed to address several key questions regarding data congruency.

First, the distinct methods of cell model construction using either genome editing (GE), or ectopic overexpression (OE) may introduce potential artifacts. We compared transcriptomic regulation in 12 ectopic overexpression and 32 genome-edited *ESR1* mutant models to their wild-type control counterparts in the absence of estrogen stimulation. Surprisingly, only a weak correlation (R=0.19) was observed in regulatory percentile correlation, highlighting significant inter-model discrepancies (Fig. 5A). Subsequently, we identified 539 and 133 genes that were more prevalently regulated in GE and OE models, respectively (delta regulatory PCT>70%), such as *BFSP2* and *ENTPD8* (Supplementary Table S2 and Supplementary Fig. S6A). Interestingly, 32 out of 133 (24%) OE-specific targets belonged to the category of noncoding RNA such as lncRNA, ncRNA, and miRNA (e.g., *LINC00886*, Supplementary Fig. S6B), suggesting that the overexpression step might particularly influence RNA processing. Importantly, this elevation was consistent in both transient doxycycline-inducible models and long-term stable overexpression models (Supplementary Fig. S6B). Pathway analysis identified a strong enrichment of innate immune response pathways (Interferon, complement, and TNFα/NFkB) in GE unique genes, possibly as a result of genome disruption. On the other hand, OE unique genes were enriched for estrogen response signatures, likely due to the overexpression of receptor levels amplifying ER-related differences (Fig. 5B). Given the differences identified, we examined how each type of model compared to clinical samples harboring *ESR1* mutations. We harmonized four published cohorts comprising 313 ER+ metastatic breast cancer patients, consisting of 246 and 67 patients harboring WT and mutant *ESR1*, respectively. Logistic regression modeling, using genes with average regulatory PCT above 60% in each model type, revealed that signatures from both models could effectively distinguish *ESR1* mutant samples from WT counterparts, with genome-edited models showing a greater performance in terms of specificity and sensitivity trade-off (Fig. 5C), suggesting a better clinical representation.

**Figure 5.**
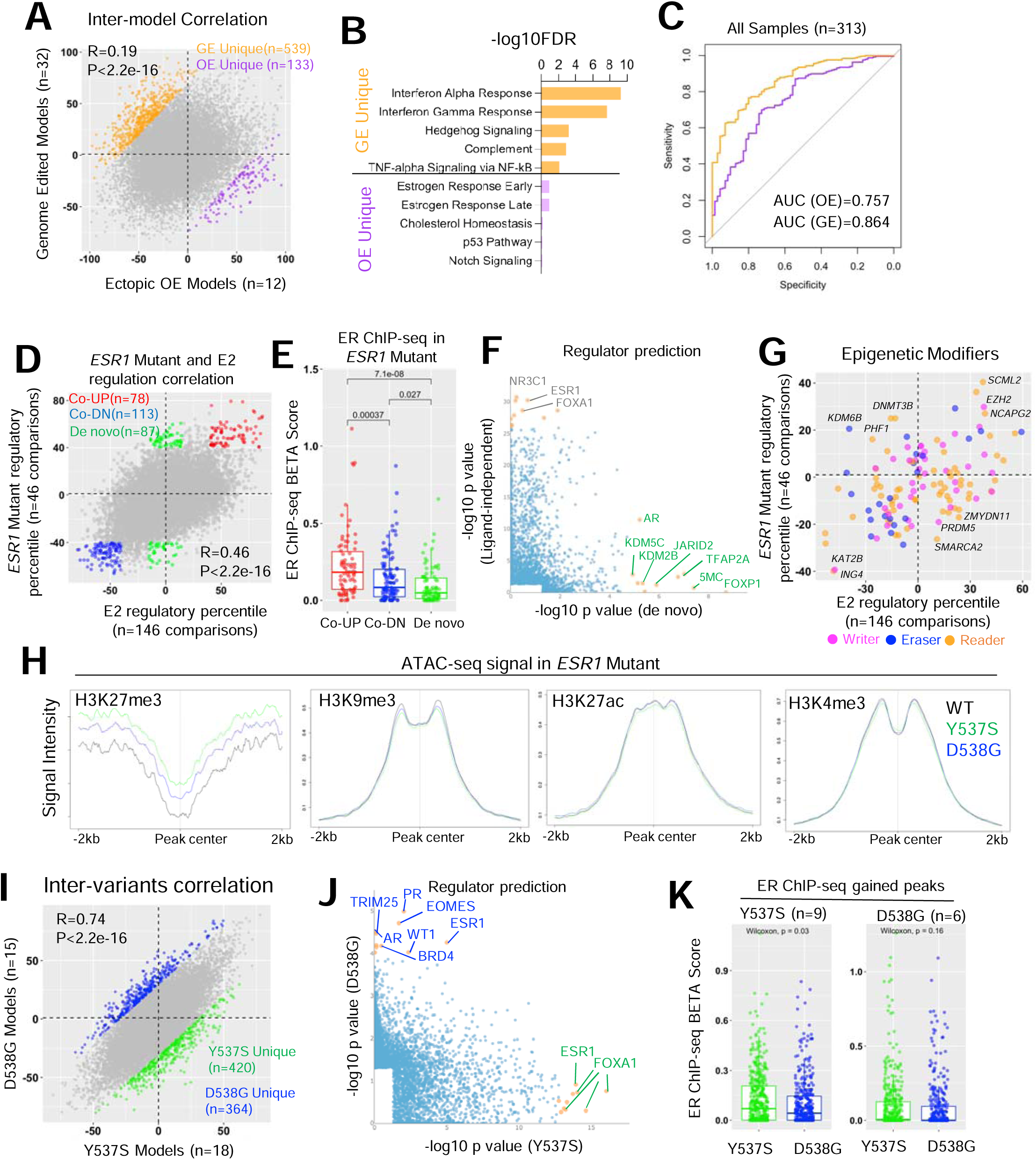
Concordance among ESR1 mutant breast cancer cell models. A. Scatter plot showing Pearson correlation of average regulatory percentile between 12 ectopic overexpression models (OE) and 32 genome-edited (GE) ESR1 mutant models of all the genes. Genes that more pronouncedly regulated by each model were highlighted (|delta regulatory percentile| >100 between the two models). B. Bar graph showing the significantly enriched Hallmark pathways in GE and OE-preferentially regulated genes in I. C. Receiver operating characteristic curve depicting the performance of GE and OE upregulated genes (average percentile >60%) as signatures in distinguishing ESR1 mutant from WT clinical samples from four merged cohort of 313 samples. D. Scatter plot showing Pearson correlation of average regulatory percentile between ESR1 mutant and estrogen treatment of all the genes. Genes are subgrouped into three parts: ligand-dependent genes (average percentile above 50% or below -50% in both conditions, red and blue) and de novo genes (average percentile above 50% in TamR and within -15%-15% in E2 treatment, green). E. Box plot representing the BETA score comparison of each selected gene subgroups in A. BETA score were calculated based on gained peaks of ESR1 mutant cells merged from 16 ChIP-seq samples normalized to their corresponding controls. Mann Whitney U test was used. F. Scatter plot showing the correlation of -log10 p values of LISA predicted regulators from ligand-dependent and de novo genes in A. Only significantly enriched regulators were shown, and top targets skewed to each side were labelled. G. Scatter plot showing correlation of average regulatory percentile between ESR1 mutant and estrogen treatment of 127 epigenetic modifiers. Targets consistently altered in both conditions or uniquely altered in one of the conditions were labelled. H. Linge plot representing the intensity of ATAC-seq signals from MCF7 WT and ESR1 mutant cell lines on different histone modification regions from ChIP-seq data of MCF7 cell line. I. Scatter plot showing Pearson correlation of average regulatory percentile between Y537S and D538G ESR1 mutant variants of all the genes. Genes that more pronouncedly regulated by Y537S and D538G were highlighted (|delta regulatory percentile| >70 between the two variants). J. Scatter plot showing the correlation of -log10 p values of LISA predicted regulators from Y537S and D538G-preferrentially regulated genes in F. Only significantly enriched regulators were shown and top targets skewed to each side were labelled. K. Box plot representing the BETA score comparison of each selected gene subgroups in F. BETA score were calculated based on gained peaks of ESR1 mutant cells merged from 9 Y537S and 6 D538G ChIP-seq samples normalized to their corresponding controls. Mann Whitney U test was used.

Multiple research teams, including ours, have reported *de novo* transcriptomic reprogramming by *ESR1* mutations, which may confer neomorphic gain-of-function phenotypes not observed by ligand-activated wild-type ER (15,64,73). However, there are considerable differences in results from different groups. We observed a strong positive correlation (R=0.46), between genes regulated in 146 estrogen-treated experiments and 46 experiments in *ESR1*-mutant cells compared to their corresponding WT controls in the absence of E2 (Fig. 5D). However, we noted 78 genes showing ligand-independent activation and 113 genes showing ligand-independent repression (average PCT>40% in both E2 and ESR1 mutation regulation) (Fig. 5D and Supplementary Table S2), such as *EGR3* and *FNBP1L*, as well as 87 de novo targets (average PCT>40% in ESR1 mutation and -15%-15% in E2 regulation), such as *CNTFR* and *CRLF1* (Supplementary Fig. S6C,S6D). Next, we investigated whether these ligand-independent and *de novo* targets are associated with mutant ER genomic binding. We analyzed gained peak sets from 16 ER ChIP-seq datasets in *ESR1*-mutant models normalized to their corresponding wild-type controls from four independent studies (32,49,52,64,74), and calculated the proximal gene regulatory score (-100 to +100 kb range of the TSS) from the peak union using Binding and Expression Target Analysis (BETA)(60). *De novo* genes showed significantly lower BETA scores compared to both upregulated and downregulated ligand-independent genes (Fig. 5E), suggesting these targets are less likely to be direct ER transcriptional targets. Utilizing the LISA algorithm (40), we predicted potential regulators of these genes. Canonical ligand-independent genes were closely associated with ER and FOXA1, while numerous epigenetic regulators at the DNA and histone levels were linked to *de novo* genes, including DNA methylation and histone modifiers such as KDM5C, JARID2, and KDM2B (Fig. 5F). Furthermore, we comprehensively assessed the regulatory percentile correlations of 127 epigenetic modifiers in *ESR1*-mutant and E2-stimulated conditions (Fig. 5G). We found that top mutant-specific mediators predominantly regulate histone H3 lysine 27 (H3K27) methylation, including the unique increase of H3K27me3 demethylase KDM6B. In line with this, we found H3K27me3 marked regions in *ESR1* mutant cells presented higher degree of chromatin accessibility by projecting ATAC-seq signals from a previous study (64), which was not discerned in other histone marked regions such as H3K9me3, H3K27ac and H3K4me3 (Fig. 5H). In summary, these analyses provide valuable insights indicating that *ESR1*-mutant cells exhibit an ER binding-independent gene expression profile via H3K27me3 modification-driven epigenetic remodeling.

Y537S and D538G are the two most frequently detected *ESR1* mutations in metastatic breast cancers resistant to endocrine therapy (49). While both mutations contribute to constitutive ER activation, pre-clinical studies have also revealed allele-specific effects (15,49,64,73), although with limited consistency across studies. We correlated the global gene regulatory percentiles and hallmark pathway alterations from 18 Y537S and 15 D538G cell model comparisons. While we observed a strong positive correlation (R=0.74), we identified 420 and 364 genes that were more prevalently regulated in Y537S and D538G variants compared to WT in the absence of E2 (Fig. 5I and Supplementary Table S2), respectively, such as *SYT3* and *SH3TC1* (Supplementary Fig. S6E). At the pathway level, Y537S was enriched for genes encoding proteins associated with metabolic functions such as glycolysis, oxidative phosphorylation, and fatty acid metabolism, while D538G regulated genes were more associated with hedgehog signaling. However, estrogen response and E2F signatures were highly upregulated in both variants (Supplementary Fig. S6F). LISA analysis revealed that Y537S-specific genes were consistently linked to ESR1 and FOXA1 across different studies, whereas the top D538G regulomes involved a diverse range of developmental-related regulators such as EOMES and WT1, as well as alternative nuclear receptors such as AR and PR (Fig. 5J). Integration of ER ChIP-seq data from 9 Y537S and 6 D538G specific models revealed that Y537S-specific genes showed significantly higher association with Y537S ER genomic binding, while such a difference was not observed in the D538G ER ChIP-seq profile (Fig. 5K). In conclusion, we found highly consistent transcriptomic regulation between ESR1 Y537S and D538G variants, while each variant possesses unique features. The unique alteration in Y537S is likely due to its distinct ER genomic binding, while D538G may involve the hijacking of nonconventional transcriptional factors to partially reshape its unique transcriptome.

### Consensus targets in *ESR1*-mutant breast cancer cell models

We have determined that despite the differences in gene expression in the *ESR1* mutant cell models, they show similarities to the transcriptomes identified in human clinical samples. With this in mind, we merged the *ESR1*-mutant cell model data with profiles of clinical specimens to pinpoint consistently regulated genes that may have been overlooked in previous studies using single models and datasets. We focused primarily on upregulated genes, given their potential utility as therapeutic targets. Among all 46 comparisons in *ESR1* mutant models, we identified 25 genes that consistently ranked among the top 10% of upregulated targets in at least 50% of comparisons. Notably, twelve of these genes also overlapped with differentially upregulated genes (padj < 0.1, log2FC > 0) detected in ESR1 mutant versus WT clinical samples from at least one of four individual metastatic breast cancer cohorts (Fig. 6A and Supplementary Fig. S7A). Surprisingly, many of these genes have not been previously studied in the context of *ESR1* mutation or even estrogen regulation, such as *RBM24* and *C5AR2*. Examination of their estrogen regulatory potential revealed that the majority of these targets are regulated via ligand-independent ER activation, while a few, such as *RND2* and *NUPR1*, are subject to *de novo* regulation (Fig. 6B). This finding underscores the potential value of comprehensive data analysis in uncovering novel insights that might otherwise be missed in single-dataset analyses.

**Figure 6.**
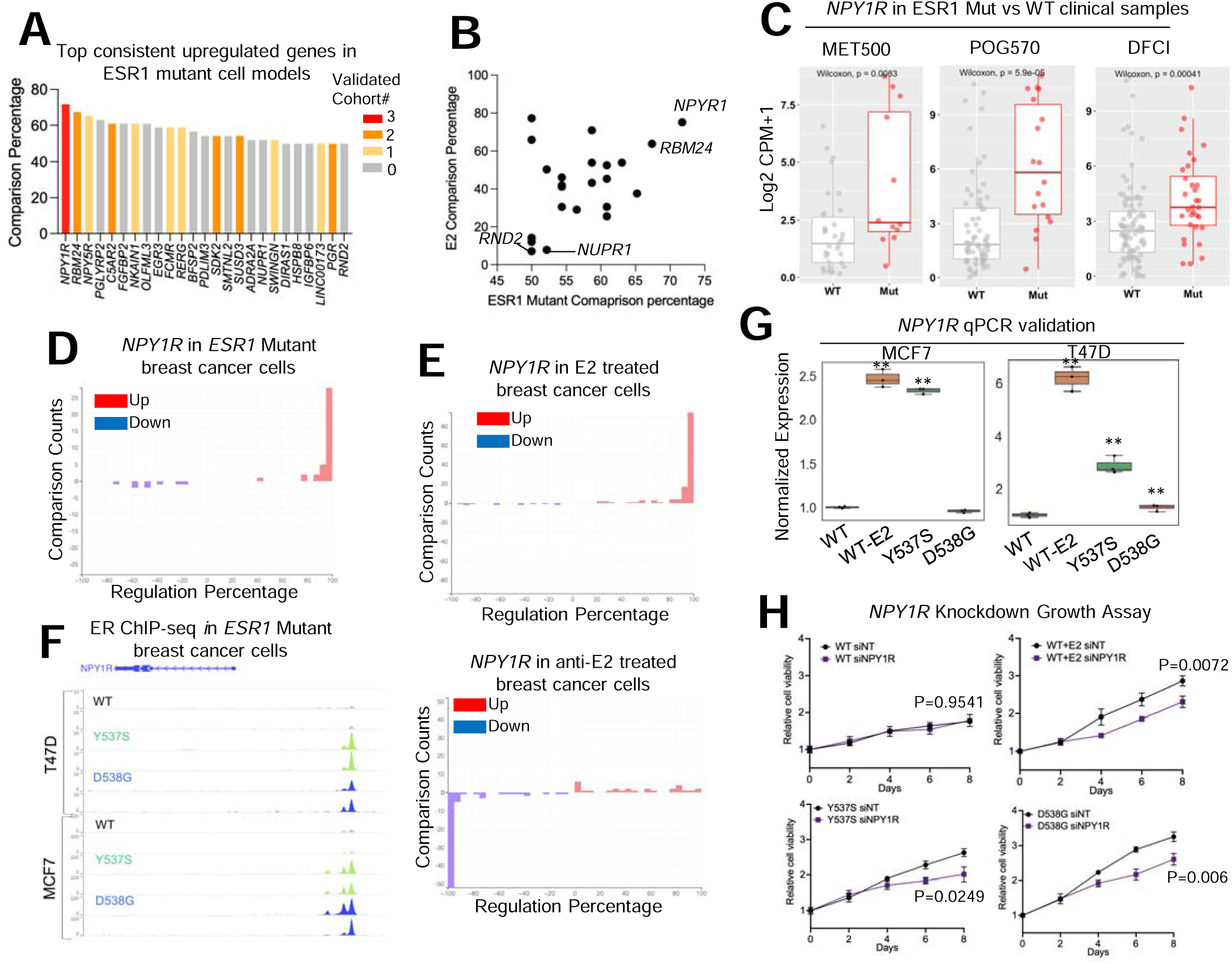
Consensus targets in ESR1 mutant breast cancer cell models. A. A bar plot ranking the genes by consisteny of falling into top10% upregulated targets in ESR1 mutant cell models. Only genes showing equal to or above 50% consistency out of 46 comparisons were shown. Color code indicates the number of cohorts (four in total) in which this gene is also identified as a significantly upregulated target in ESR1 mutant samples. B. Scatter plot showing the correlation of average regulatory percentile between ESR1 mutant and estrogen treatment of 25 targets in A. C. Box plots showing the expression of NPY1R between ESR1 WT and mutant samples from the three cohorts. D and E. Bar plot showing the distribution of regulatory percentile of NPY1R in all the comparisons from ESR1 mutant cells (D), estrogen and ER modulator treatments (E) experiments. Percentiles are ranged between -100 to +100 and larger number indicates stronger regulation. F. Genomic track view of ER ChIP-seq signal from ESR1 WT and ESR1 mutant models in two cell lines from GSE148277. Genomic track scale is adjusted to the identical level for all the samples. G. Box plots showing the relative expression of NPY1R in genome-edited MCF7 and T47D ESR1 mutant cells and WT treated with E2 condition. Δ ΔCt method is used and p values were calculated using one-way ANOVA. H. Line plot showing the growth rate of T47D WT and ESR1 mutant cells in the presence of scramble (siNT) and NPY1R siRNA transfection for 8 days. Two-way ANOVA was used for statistic comparison. Experiment has been repeated for 3 times.

Of particular interest, we observed two neuropeptide Y receptor (NPYR) genes, *NPY1R* and *NPY5R*, among the top three consistently upregulated targets in all the *ESR1* mutant models. NPY1R has been studied in the context of endocrine sensitivity in ER+ breast cancer(76) although it specific role(s) in *ESR1* mutant breast cancers has not been explored. Initially, we confirmed a significant upregulation of *NPY1R* in *ESR1* mutant tumors in three out of four clinical cohorts (Fig. 6C), along with its consistent elevation in *ESR1* mutant cell models (irrespective of cell line and *ESR1* mutant type) (Fig. 6D and Supplementary Fig. S7B). Notably, *NPY1R* demonstrated dominant upregulation upon E2 stimulation, while its expression was suppressed by inhibitors of ER signaling (Fig. 6E). Furthermore, we consistently observed gained ER binding at the proximal region of the *NPY1R* locus in *ESR1* mutant cells (Fig. 6F). Taken together, these findings identify *NPY1R* as one of the most consistent and universal upregulated targets in *ESR1* mutant breast cancer, likely driven by the constitutive activation of mutant ER and their increased binding activity at this gene locus. To further investigate its role in endocrine resistance in ESR1 mutant cells, we conducted knockdown experiments using MCF7 and T47D genome-edited cell models(73). qRT-PCR validated its elevated expression in both T47D cell lines and MCF7 Y537S models (Fig. 6G). Remarkably, knockdown of *NPY1R* significantly inhibited the growth of T47D WT cells under E2 stimulation, as well as the ligand-independent growth of *ESR1* mutant cells (Fig. 6H and Supplementary Fig. S7C), underscoring the essential role of NPY1R in estrogen-dependent and *ESR1* mutant breast cancer cell growth.

## Discussion

The increase in multiomic cancer data represents an invaluable resource, yet integration and cross-comparison between individual studies poses significant challenges. The initial EstroGene database (35) laid a solid foundation by providing a unified analysis of multiple datasets and offering a versatile tool for comprehensive exploration of transcriptional responses to estrogens in validated models of breast cancer. EstroGene2.0 represents a substantial enhancement by incorporating experimental data on responses to endocrine response and on resistance to various interventions. EstroGene2.0 has a robust and user-friendly interface, empowering researchers to assess individual genes or gene signatures generated under specific experimental conditions. With access to both gene expression data and ER proximity binding information, users can investigate cross-dataset consistency. EstroGene2.0 provides an unprecedented opportunity to dissect the technical and biological intricacies inherent in response to endocrine therapies.

In this study, we identified a more heterogeneous transcriptomic response in tamoxifen treatment compared to fulvestrant. This disparity may stem from the fact that tamoxifen is a partial ER agonist (77). For instance, the diverse transcriptomic reprogramming triggered by different doses of tamoxifen may arise from the recruitment of distinct co-activators and the distinct status of monomeric versus dimeric ER. Moreover, despite an overall positive correlation between gene regulation across all ER-modulators evaluated, each compound type can regulate distinct gene clusters. Notably, the ER LDD ARV-471 exhibited similar effects to SERDs, attributable to their shared mechanism of ER downregulation. Among SERMs, tamoxifen, raloxifene, and bazedoxifene elicited markedly different effects compared to lasofoxifene and endoxifen. While this variation may reflect the limited experiments included for the latter two compounds and the differential distribution of cell model and modality used, it is plausible that different SERMs confer unique molecular portraits, warranting further investigation. It is also possible that these agents may differentially regulates other estrogen receptors such as ERβ and G Protein-Coupled Estrogen Receptor (GPER). Furthermore, we observed a tight link between transcriptomic response and cell line context. Integration with RIME data revealed the differential dependency of cell-line-specific transcriptional regulators. For instance, ER demonstrated a stronger binding affinity with co-repressor TLE3 in the CAMA1 cell line, and ER modulator treatment could more profoundly reshape TLE3/ER-related downstream targets, resulting in a distinct molecular signature compared to other cell lines. Importantly, the identification of these context-dependent effects underscores the necessity for future studies to encompass a wider spectrum of models when testing ER modulators. The examination of more clinically relevant models, such as patient-derived organoids and immunocompetent animal models, is warranted to further elucidate the complex interplay between ER modulators and complexed breast cancer biology.

Endocrine resistance presents a complex landscape of molecular determinants, as our integrated analysis revealed. We observed interferon response signaling showing as a consistent negative association with estrogen signaling shift. Mutual regulation between these two axes has been previously reported in wild-type breast cancer cells. Specifically, activation of ER signaling can inhibit type I interferon response by restricting the engagement of the IFN-stimulated gene factor 3 (ISGF3) complex at the ISG promoters and disrupting the ISGF3 complex via interaction with STAT2 (78). Additionally, it has been demonstrated that the addition of type I and II interferon treatment can enhance the anti-proliferative effects of tamoxifen in breast cancer cell lines (79). These findings suggest that the plasticity of interferon response may buffer the alterations in estrogen signaling seen in endocrine-resistant settings, either boosting ER signaling in ER repressive models like LTED or attenuating ER signaling in ER hyperactivation models like ESR1 mutants. Combining therapies that increase interferon signaling with standard-of-care ER modulator treatment may offer a potential avenue to overcome this resistance. Analysis from the POETIC clinical trial (47) also highlights a potential prognostic role of interferon response in endocrine-resistant cases. However, caution should be exercised when interpreting these data, as the interferon response signature from bulk tumor sequencing may also be influenced by immune infiltration patterns. Additionally, the source of divergent interferon response in these models warrants further investigation. One possibility relates to the method of model construction; for instance, tamoxifen is known to induce DNA damage (80), which could trigger innate immune responses via the cGAS-STING pathway. Therefore, the varying doses and durations of tamoxifen selection in various studies could introduce different degrees of interferon response, and some selected clones may harbor mitigated interferon signaling. It once again emphasizes the importance of incorporating optimal control into experimental design, particularly for models undergone extensive engineering or selection.

*ESR1* mutations are likely drivers of resistance to endocrine therapy and are associated with metastatic disease. While pre-clinical cell models have extensively probed this phenomenon, the consistency of findings from in vitro models remains to be examined. In our prior studies we uncovered epigenetic dysregulation in *ESR1* mutant cells, notably the pronounced enrichment of FOXA1, CTCF, and OCT1 in open chromatin present in *ESR1* mutant cells (64), alongside alterations in chromatin interaction patterns driven by CTCF/cohesin complex remodeling(52). Building upon this foundation, investigation of 46 *ESR1* mutant transcriptomic profiles highlights a tight association between epigenetic regulators and *de novo* transcriptomes. This correlation may stem from increased accessibility at facultative heterochromatin regions marked by H3K27me3, while changes at H3K27ac-marked regions were not discerned. These insights underscore how *ESR1* mutations, by reducing repressive effects at existing heterochromatin marks, can sufficiently reshape the transcriptome, and the elevation of H3K27me3 demethylase KDM6A in *ESR1* mutant cells supports this proposition. Furthermore, our observations point towards the involvement of DNA methylation in regulating these de novo genes, aligning with findings of mutant-specific alterations in DNA methyltransferase, such as *DNMT3B*. Notably, previous pharmacological studies have revealed heightened sensitivity of *ESR1* mutant cell models towards epigenetic modifier inhibitors like OTX015, which targets the bromodomain and extra terminal domain (BET)(81). These findings underscore the potential for further exploration into the mechanisms of epigenetic reprogramming in ESR1 mutant breast cancer, offering a promising avenue for uncovering novel vulnerabilities and therapeutic strategies.

Previous preclinical studies have shed light on allele-specific functions of *ESR1* Y537S and D538G variants, an idea bolstered by several clinical observations. For example, the BOLERO2 trial showcased differences in overall survival and response to everolimus treatment between tumors harboring Y537S and D538G mutations(82). Our comprehensive analysis, comparing 18 Y537S and 15 D538G-induced transcriptomic experiments, reveals an overall high degree of similarity but we also note unique biology attributable to each mutant. Specifically, our findings suggest that Y537S variant-specific transcriptomes are predominantly regulated by ER genomic binding, whereas the shifts unique to D538G variants stem from a diverse array of ER-independent functions. This aligns with previous clinical data analyses indicating that Y537S mutant tumors exhibit a more pronounced enhancement of ER signaling(49). Given that these models predominantly utilize MCF7 and T47D cell lines, there is an urgent need to establish additional *ESR1*-mutant cell lines and PDX and organoid models with diverse genetic backgrounds to corroborate these observations. Lastly, leveraging our comprehensive data collection, we identified several consistently upregulated genes in both *ESR1* mutant models and clinical samples, many of which have not been previously described. Notably, two neuropeptide Y receptor (NPYR) genes, *NPY1R* and NPY5R, emerged among the top three genes in our analysis. Our additional experiments validated the increased expression of *NPY1R* in *ESR1* mutant cells and its role in conferring growth dependency. Interestingly, a recent study also identified NPY1R as an estrogen (E2)-activated gene in breast cancer cell lines(76), and showed that activation of NPY1R by ligand stimulation partially inhibited E2-driven cell proliferation. This finding contrasts with our finding that knockdown of *NPY1R* reduced estrogen stimulated and *ESR1* mutant mediated growth, however, our studies were performed in the absence of *NPY1R* ligand activation. It is plausible that the inactive state of *NPY1R* itself may engage in crosstalk with other signaling pathways at baseline, contributing to the ligand-independent growth phenotype in *ESR1* mutant cells, given previous studies demonstrating its extensive crosstalk network in central nervous system diseases(83).

In conclusion, the EstroGene2.0 database stands as a user-friendly platform tailored for the analysis and visualization of endocrine response and resistance in breast cancer. Similar to our initial observations stemming from Estrogene1.0, meta-analysis of multiple experiments and model systems reveals numerous novel findings not possible from single model studies. Our future vision extends beyond its current capabilities, as we aim to enrich the platform with additional epigenetic datasets, including histone modifications (e.g., histone ChIP-seq) and chromatin interaction profiles (e.g., Hi-C), thereby providing deeper mechanistic insights in the future. Furthermore, our roadmap includes the integration of datasets from clinical trials involving various endocrine therapies in ER+ breast cancer. This expansion will culminate in the development of a consensus network informed by user input from clinical settings, facilitating informed decision-making in patient care. Moreover, our studies have generated a plethora of hypotheses, underscoring the need for future experimental validation and analysis of clinical specimens. We anticipate that the EstroGene2.0 database will serve as a catalyst in the quest to overcome resistance to endocrine therapy in breast cancer patients and beyond, ultimately contributing to improved patient outcomes.

## Data Availability

Details of all the curated all data sets used for database construction are summarized in Supplementary Table S1. This includes all the associated publication information, GEO accession numbers, experimental designs including cell models, compound dose, duration, resistant model generation methods, library preparation method and NGS sequencing platforms. For ChIP-seq integration with other epigenetic data, preprocessed BED files for ChIP-seq of H3K27ac, H3K9me3 and H3K27me3 in T47D were downloaded from GSE63109(63). T47D ESR1 mutant cell ATAC-seq data were downloaded from GSE148277(64). For clinical cohort analysis, expression data and sample metadata of POETIC trial were downloaded from GSE105777(47). For *ESR1* mutant metastatic breast cancer cohorts, data set availability was described in our previous publication(49), including our local WCRC cohort (46 *ESR1* WT and eight mutant tumors) and three previously reported cohorts—MET500(50) (34 *ESR1* WT and 12 mutant tumors), POG570(51) (68 *ESR1* WT and 18 mutant tumors), and Dana-Farber Cancer Institute (DFCI; 98 *ESR1* WT and 32 mutant tumors)(52).

## Acknowledgements

The authors would like to thank Dr. Ciara Metcalfe (Genetech, Inc.) for granting early access of GEO deposition from their study to allow integrative meta-analysis in this study.

