## Supplementary Figures for "EstroGene2.0: A multi-omic database of response to estrogens, ER-modulators, and resistance to endocrine therapies in breast cancer"

### Supplementary Figure S1

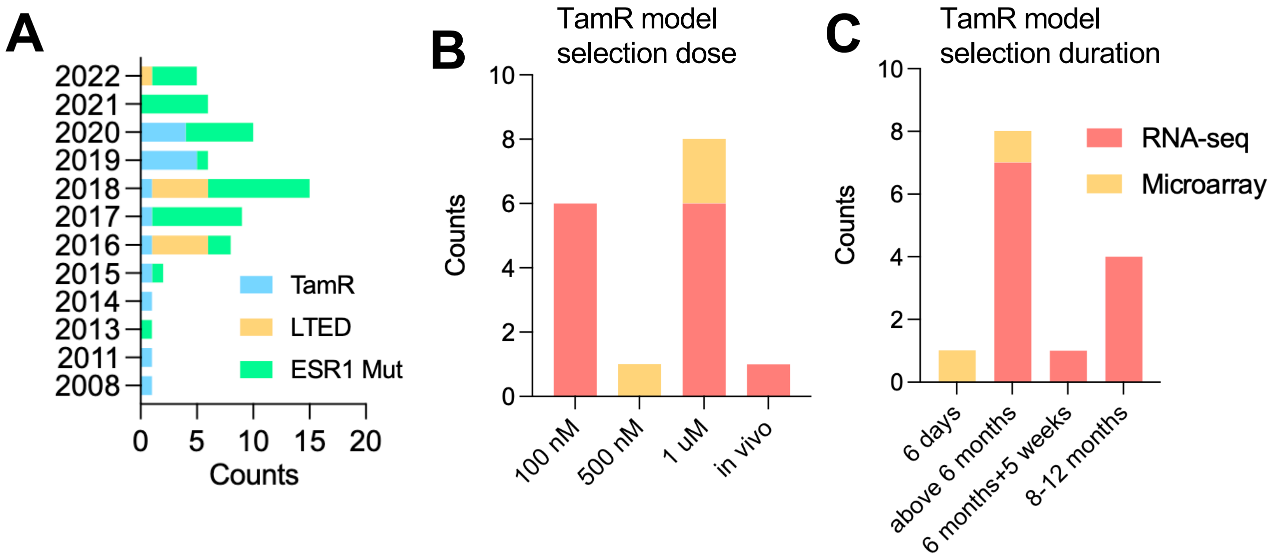


Supplementary Figure S1 (related to Fig. 1)

A. Stacked histogram showing the metadata separated by technologies and three endocrine resistant model types.

B and C. Stacked histogram summarizing the dose (B) and duration (C) of tamoxifen resistant model selection separated by technologies.

### Supplementary Figure S2

**A**

B-E

F-H

**Modality**

Multi-select

- ER ChIP-seq
- Microarray
- RNA-seq

**Cell Line**

Multi-select

- MCF7
- MDAMB453
- SUM44
- SUM44PE
- T47D

**Cell Model Type**

Multi-select

- AAV Genome-edited
- CRISPR/Cas9
- Dox-inducible Overexpression
- Naturally Occurred in LTED condition

| Data Set ID | Experiment ID | Modality | Cell model | Cell Line | Replicates | Cell Model Type | Variant Type | Ins |
|---|---|---|---|---|---|---|---|---|
| ERR7 | 1 MUTR1_1 | RNA-seq | ESR1 Mutation | T47D | 4 | CRISPR/Cas9 | Y537S, D538G | University |
| ERR7 | MUTR1_2 | RNA-seq | ESR1 Mutation | MCF7 | 4 | AAV Genome-edited | Y537S, D538G | University |
| ERR8 | MUTR2 | RNA-seq | ESR1 Mutation | MCF7 | 3 | CRISPR/Cas9 | Y537S | Imperial C |
| ERR9 | MUTR3 | RNA-seq | ESR1 Mutation | T47D | 3 | CRISPR/Cas9 | Y537S, D538G | University |
| ERR10 | MUTR4_1 | RNA-seq | ESR1 Mutation | T47D | 3 | Dox-inducible Overex... | Y537S | Dana-Fart |
| ERR10 | MUTR4_2 | RNA-seq | ESR1 Mutation | MCF7 | 3 | Dox-inducible Overex... | Y537S, D538G, Y537N | Dana-Fart |
| ERR11 | MUTR5 | RNA-seq | ESR1 Mutation | MCF7 | 6 | CRISPR/Cas9 | Y537S, D538G, L536... | Imperial C |
| ERR12 | MUTR6 | RNA-seq | ESR1 Mutation | MCF7 | 2 | CRISPR/Cas9 | Y537S, D538G | UT Southw |
| ERR13 | MUTR7_1 | RNA-seq | ESR1 Mutation | MCF7 | 2 | CRISPR/Cas9 | Y537S, D538G | University |
| ERR13 | MUTR7_2 | RNA-seq | ESR1 Mutation | T47D | 2 | CRISPR/Cas9 | Y537S, D538G | University |
| ERR15 | MUTR8_1 | RNA-seq | ESR1 Mutation/LTED | MCF7 | 3 | Naturally Occurred in ... | Y537C | Institute o |
| ERR15 | MUTR8_2 | RNA-seq | ESR1 Mutation/ITED | SUM44PE | 3 | Naturally Occurred in ... | Y537S | Institute o |
| ERR16 | MUTR9 | RNA-seq | ESR1 Mutation | T47D | 1 | CRISPR/Cas9 | Y537S, D538G | Baylor Col |
| ERR17 | MUTR10 | RNA-seq | ESR1 Mutation | MCF7 | 2 | TALEN Genome Editing | Y537S, D538G,E380Q | Dana-Fart |
| ERM4 | MUTM1 | Microarray | ESR1 Mutation | MCF7 | 3 | Stable Overexpression | Y537S, S463P, Y537... | MSKCC |
| ERM5 | MUTM2 | Microarray | ESR1 Mutation | MCF7 | 3 | Stable Overexpression | Y537S, D538G | H3 Biome |
| ERM6 | MUTM3_1 | Microarray | ESR1 Mutation | MCF7 | 3 | Stable Overexpression | Y537S | Baylor Col |
| ERM6 | MUTM3_2 | Microarray | ESR1 Mutation | ZR751 | 3 | Stable Overexpression | Y537S | Baylor Col |
| ERM7 | MUTM4 | Microarray | ESR1 Mutation | T47D | 3 | Stable Overexpression | Y537S | University |
| ERC1 | 2 MUTC1_1 | ER ChIP-seq | ESR1 Mutation | MCF7 | 1 | AAV Genome-edited | Y537S, D538G | University |
| ERC1 | MUTC1_2 | ER ChIP-seq | ESR1 Mutation | T47D | 1 | CRISPR/Cas9 | Y537S, D538G | University |
| ERC2 | MUTC2_1 | ER ChIP-seq | ESR1 Mutation | MCF7 | 2 | CRISPR/Cas9 | Y537S, D538G | University |
| ERC2 | MUTC2_2 | ER ChIP-seq | ESR1 Mutation | T47D | 2 | CRISPR/Cas9 | Y537S, D538G | University |
| ERC3 | MUTC3 | ER ChIP-seq | ESR1 Mutation | MCF7 | 1 | CRISPR/Cas9 | Y537S | Imperial C |

**B**

| GSM ID | Cell Line | Sample ID | ESR1 Genotype | GREB1 |
|---|---|---|---|---|
| GSM2392590 | T47D | MUTR1_1_DG1 | D538G | 6.710 |
| GSM2392591 | T47D | MUTR1_1_DG2 | D538G | 6.505 |
| GSM2392592 | T47D | MUTR1_1_DG3 | D538G | 6.628 |
| GSM2392593 | T47D | MUTR1_1_DG4 | D538G | 6.365 |
| GSM2392582 | T47D | MUTR1_1_WT1 | WT | 5.806 |
| GSM2392583 | T47D | MUTR1_1_WT2 | WT | 5.689 |
| GSM2392584 | T47D | MUTR1_1_WT3 | WT | 5.776 |
| GSM2392585 | T47D | MUTR1_1_WT4 | WT | 5.770 |
| GSM2392586 | T47D | MUTR1_1_YS1 | Y537S | 7.385 |
| GSM2392587 | T47D | MUTR1_1_YS2 | Y537S | 7.332 |
| GSM2392588 | T47D | MUTR1_1_YS3 | Y537S | 7.697 |
| GSM2392589 | T47D | MUTR1_1_YS4 | Y537S | 7.420 |

**C**

| Comparison | log2 Fold Change | Adjusted P-value |
|---|---|---|
| D538G over WT | 0.8002 | 3.18e-41 |
| Y537S over WT | 1.713 | 5.43e-79 |

**D**

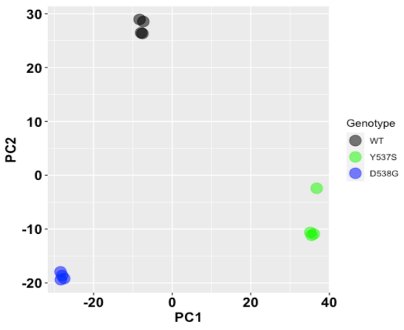


**E**

- MUTR1_1_D538G_over_WT_DEseq_output.csv
- MUTR1_1_log2CPM_Expression_Matrix.csv
- MUTR1_1_Y537S_over_WT_DEseq_output.csv

**F**

| SRR ID | Cell Line | Sample ID | ESR1 Genotype | Mapped Reads(M) | Total Peaks |
|---|---|---|---|---|---|
| SRR8444288 | MCF7 | MCF7_D538G_Vehicle | D538G | 17.36 | 399 |
| SRR8444282 | MCF7 | MCF7_WT_Vehicle | WT | 19.59 | 26 |
| SRR8444285 | MCF7 | MCF7_Y537S_Vehicle | Y537S | 18.18 | 265 |

**G**

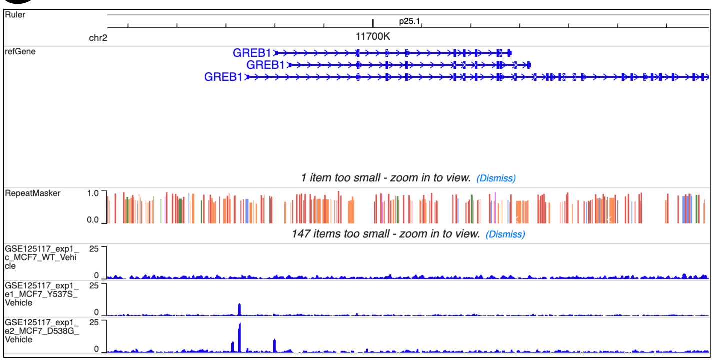


**H**

- MUTC1_1_MCF7_D538G_Vehicle_peaks.narrowPeak
- MUTC1_1_MCF7_WT_Vehicle_peaks.narrowPeak
- MUTC1_1_MCF7_Y537S_Vehicle_peaks.narrowPeak
- MUTC1_1_read.counts.rpkm.csv

### Supplementary Figure S2 (Continued)

Supplementary Figure S2. (related to Fig. 2)
A. A screen shot from EstroGene2.0 browser of “METADATA” tab of ESR1 mutant models. Red dots indicate the two example data sets elaborated in the following panels.
B-E. Screen shot from EstroGene2.0 browser of searchable “Gene Expression Matrix” panel (B), ”Principal Component Analysis” (C) and “Download” tabs for MUTR1_1 RNA-seq experiment, as an example of gene expression profiling.
F-H. Screen shot from EstroGene2.0 browser of “Experimental Metadata” (F), “Genomic Track Veiw” (G) and “Download” tabs for MUTC1_1 ER ChIP-seq experiment, as an example of ChIP-seq profiling.

### Supplementary Figure S3

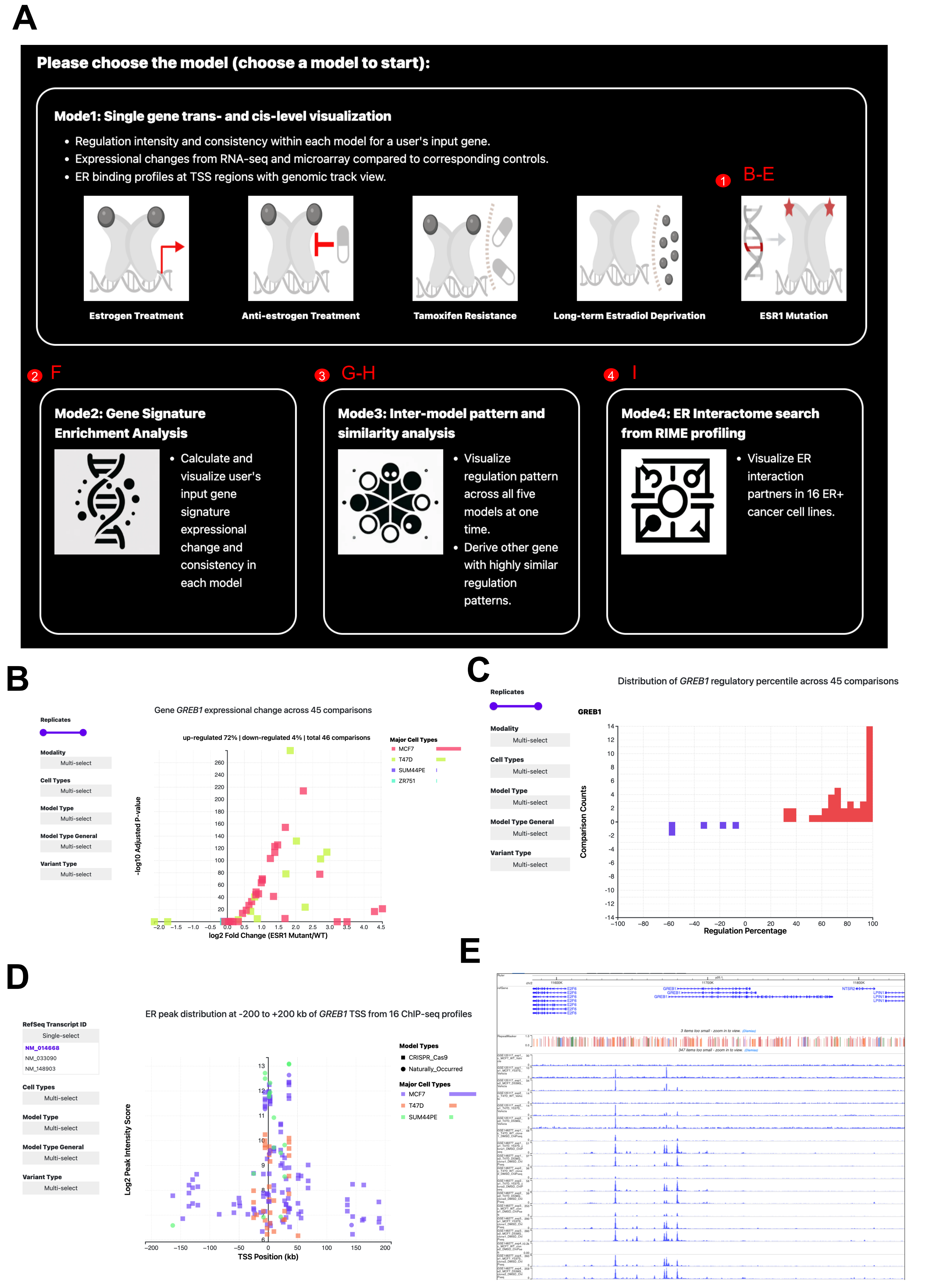

### Supplementary Figure S3 (Continued)

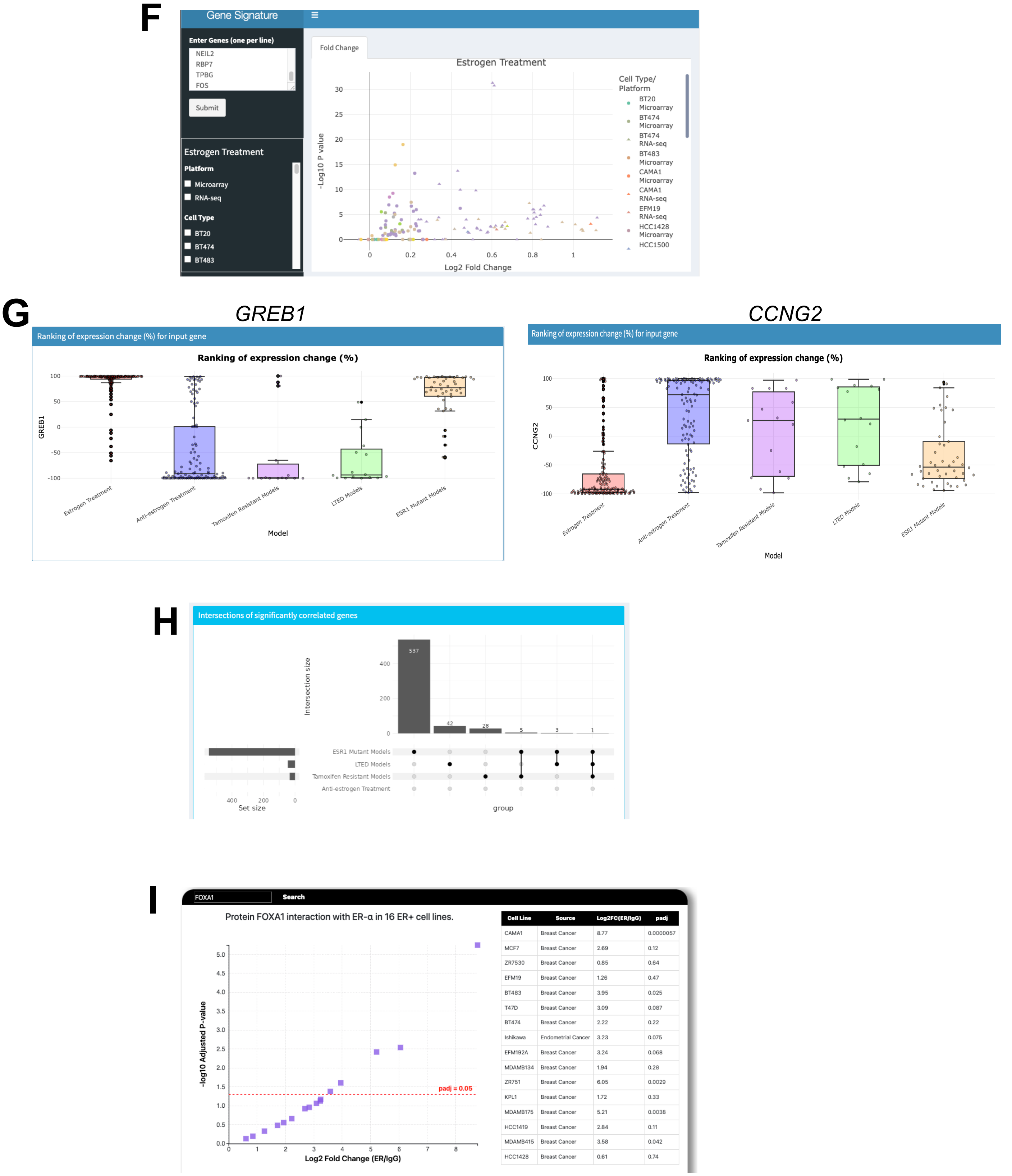


| Cell Line | Source | Log2FC(ER/IgG) | padj |
|---|---|---|---|
| CAMA1 | Breast Cancer | 8.77 | 0.0000057 |
| MCF7 | Breast Cancer | 2.69 | 0.12 |
| ZR7530 | Breast Cancer | 0.85 | 0.64 |
| EFM19 | Breast Cancer | 1.26 | 0.47 |
| BT483 | Breast Cancer | 3.95 | 0.025 |
| T47D | Breast Cancer | 3.09 | 0.087 |
| BT474 | Breast Cancer | 2.22 | 0.22 |
| Ishikawa | Endometrial Cancer | 3.23 | 0.075 |
| EFM192A | Breast Cancer | 3.24 | 0.068 |
| MDAMB134 | Breast Cancer | 1.94 | 0.28 |
| ZR751 | Breast Cancer | 6.05 | 0.0029 |
| KPL1 | Breast Cancer | 1.72 | 0.33 |
| MDAMB175 | Breast Cancer | 5.21 | 0.0038 |
| HCC1419 | Breast Cancer | 2.84 | 0.11 |
| MDAMB415 | Breast Cancer | 3.58 | 0.042 |
| HCC1428 | Breast Cancer | 0.61 | 0.74 |

Supplementary Figure S3. (related to Fig. 2)

A. A screen shot from EstroGene2.0 browser of “ANALYSIS” tab. Red dots indicate the sections elaborated in the following panels.

B-E. Screen shots from EstroGene2.0 browser of “Volcano Plot” (B) and “Percentile Plot” (C) from gene entry “GREB1” in ESR1 mutation section, and ER ChIP-seq data visualization of “TSS Region View” (D) and “Genomic Track View” (E) from from gene entry “GREB1” in ESR1 mutation section.

F. A screen shot of output from EstroGene2.0 browser of Mode2: Gene Signature Enrichment Analysis, using EstroGene meta-signature as an input and the enrichment score in estrogen treatment experiments were plotted.

G and H. Screen shots of output from EstroGene2.0 browser of Mode3: Inter-model pattern and similarity analysis, using gene GREB1 as input. Output indicates the box plot view of overall regulatory percentile of each comparisons across five different sections for *GREB1* and *CCNG2* (G) and significantly similar genes shared across all four sections (H) for *GREB1*.

I. A screen shot of output from EstroGene2.0 browser of Mode4: ER Interactome search from RIME profiling, using FOXA1 as a default input. Volcano plot depicting the log2FC and -log10padj normalized to IgG control in 16 ER+ cancer cell lines.

### Supplementary Figure S4

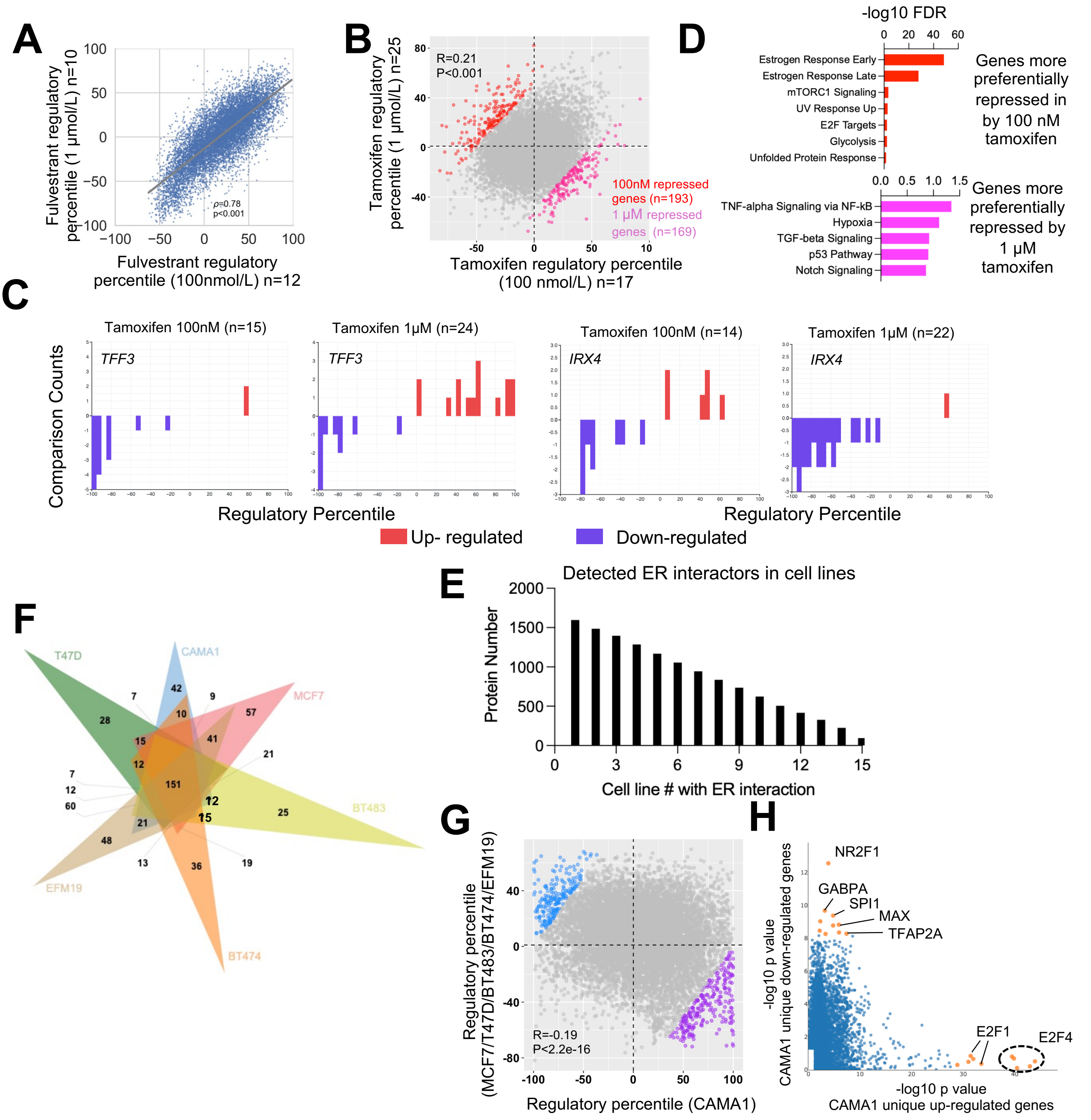


Supplementary Figure S4 (related to Fig. 3)

A and B. Scattered plots representing the Pearson correlation of regulatory percentile between 100 nM and 1 μM fulvestrant (A) and tamoxifen (B) treatment. For tamoxifen treatment experiments, genes that more pronouncedly repressed by high and low dose were highlighted in pink and red respectively. (|delta regulatory percentile| >100 between the two models).

C. Bar plot showing the distribution of regulatory percentile of TFF3 and IRX4 in all the comparisons from 100nM and 1 μM tamoxifen treatment. Percentiles are ranged between -100 to +100 and larger number indicates stronger regulation.

D. Bar graph showing the significantly enriched Hallmark pathways in high and low tamoxifen-preferentially repressed genes indicated in B.

E. Bar plot representing the detected ER interactor numbers by RIME (log2FC>5, padj<0.05 to IgG) in number of breast cancer cell lines cumulative from 1-15.

F. Venn diagram showing the overlapping of ER interactors from RIME experiment (log2FC>5, padj<0.05 to IgG) among CAMA1, MCF7, T47D, BT483, BT474 and EFM19 cells.

G. Scattered plots representing the Pearson correlation of regulatory percentile between CAMA1 and average of MCF7, T47D, BT483, BT474 and EFM19 cells under tamoxifen treatment. Genes that are differentially regulated by CAMA1 and other five cell lines were highlighted in blue and purple respectively. (|delta regulatory percentile| >100 between the two subgorups).

H. Scattered plot showing the correlation of -log10 p values of LISA predicted regulators from differentially regulated genes between CAMA1 and MCF7, T47D, BT483, BT474 and EFM19 cells in F. Only significantly enriched regulators were shown and top targets skewed to each side were labelled.

### Supplementary Figure S5

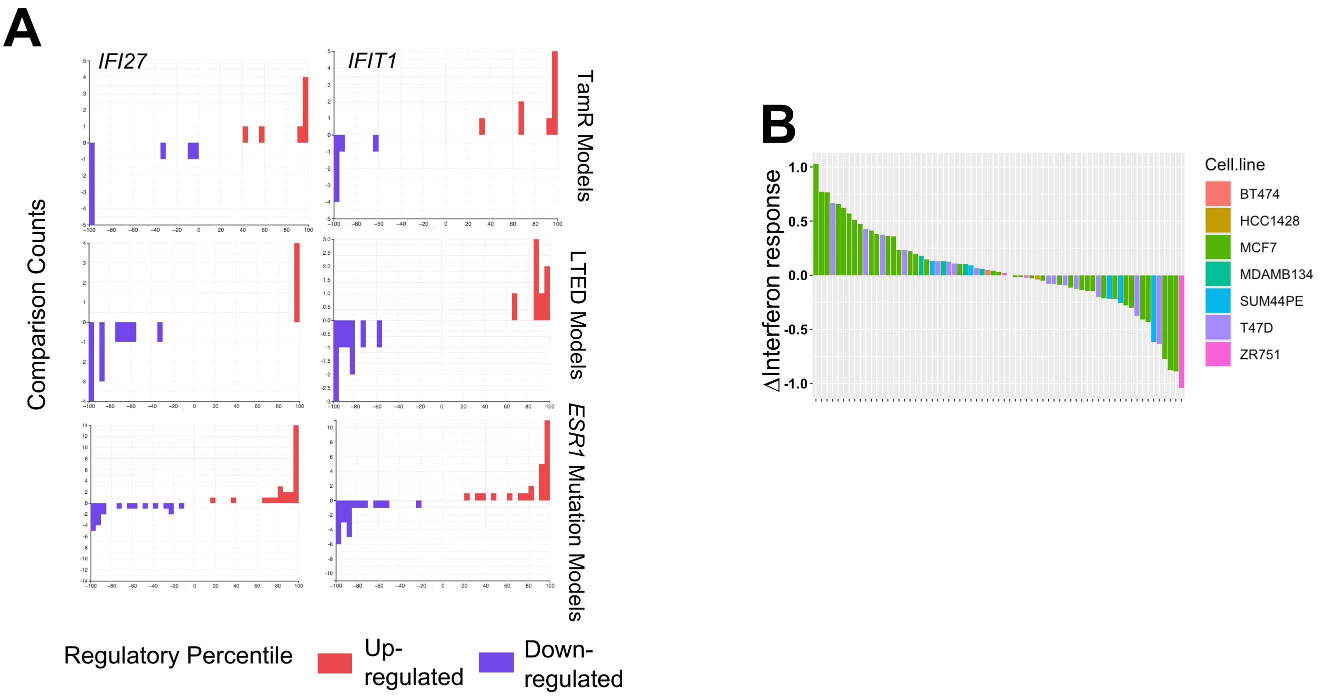


Supplementary Figure S5 (related to Fig. 4)

A. Bar plot showing the distribution of regulatory percentile of IFI27 and IFIT1 in all the comparisons from TamR, LTED and ESR1 mutations respectively. Percentiles are ranged between -100 to +100 and larger number indicates stronger regulation.

B. Bar graph showing the the alteration of GSVA enrichment scores of mean of interferon response α and γ signatures in all the comparisons from endocrine resistant models. Color code indicate specific cell lines used.

### Supplementary Figure S7

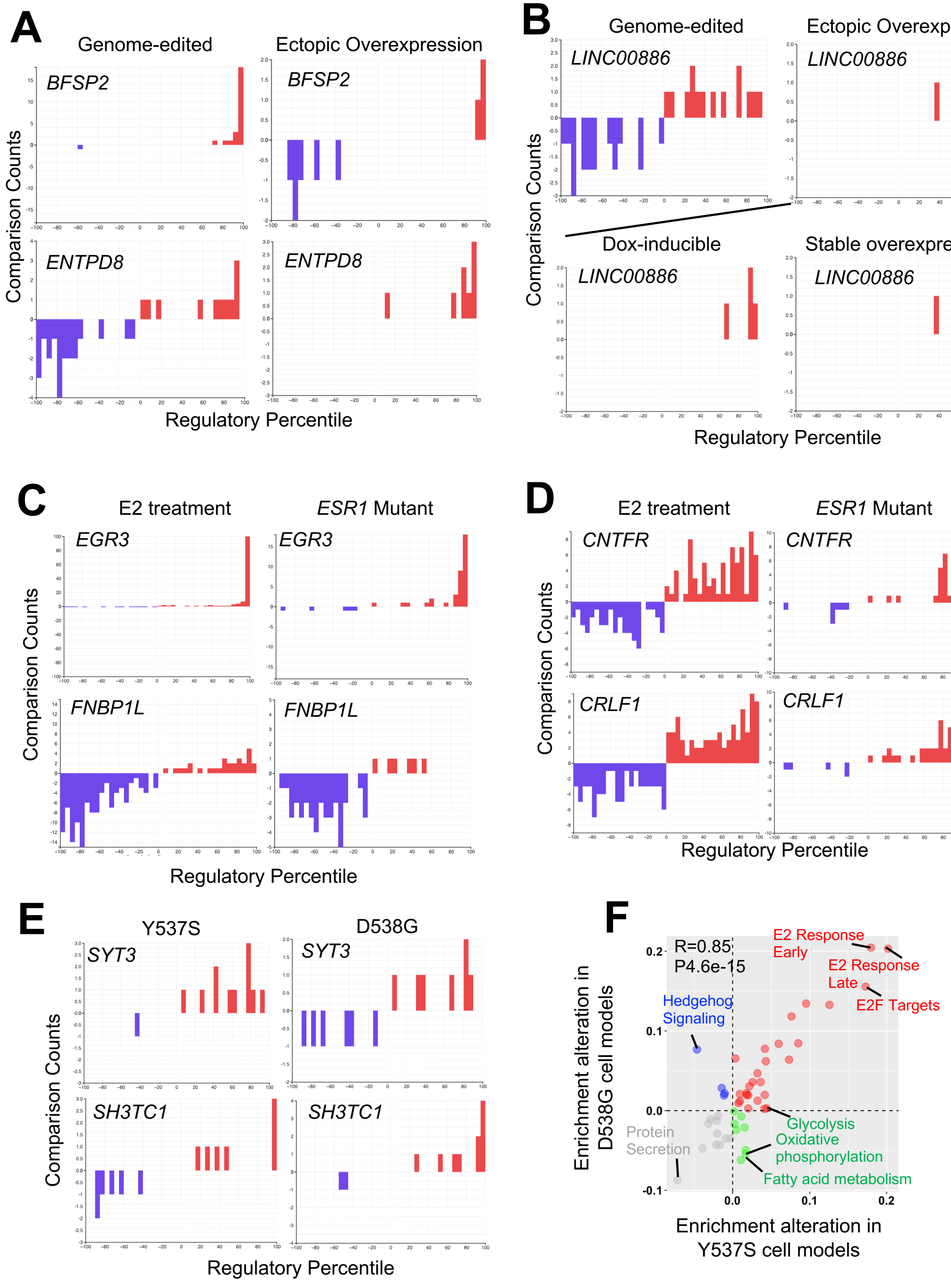


Supplementary Figure S6 (related to Fig. 5)
A-E. Bar plot showing the distribution of regulatory percentile of indicated genes in the selected comparisons from endocrine resistant cells. Percentiles are ranged between -100 to +100 and larger number indicates stronger regulation.
F. Scattered plot showing Pearson correlation between Hallmark signature enrichment score alteration in Y537S and D538G models normalized to the corresponding WT controls.

### Supplementary Figure S8

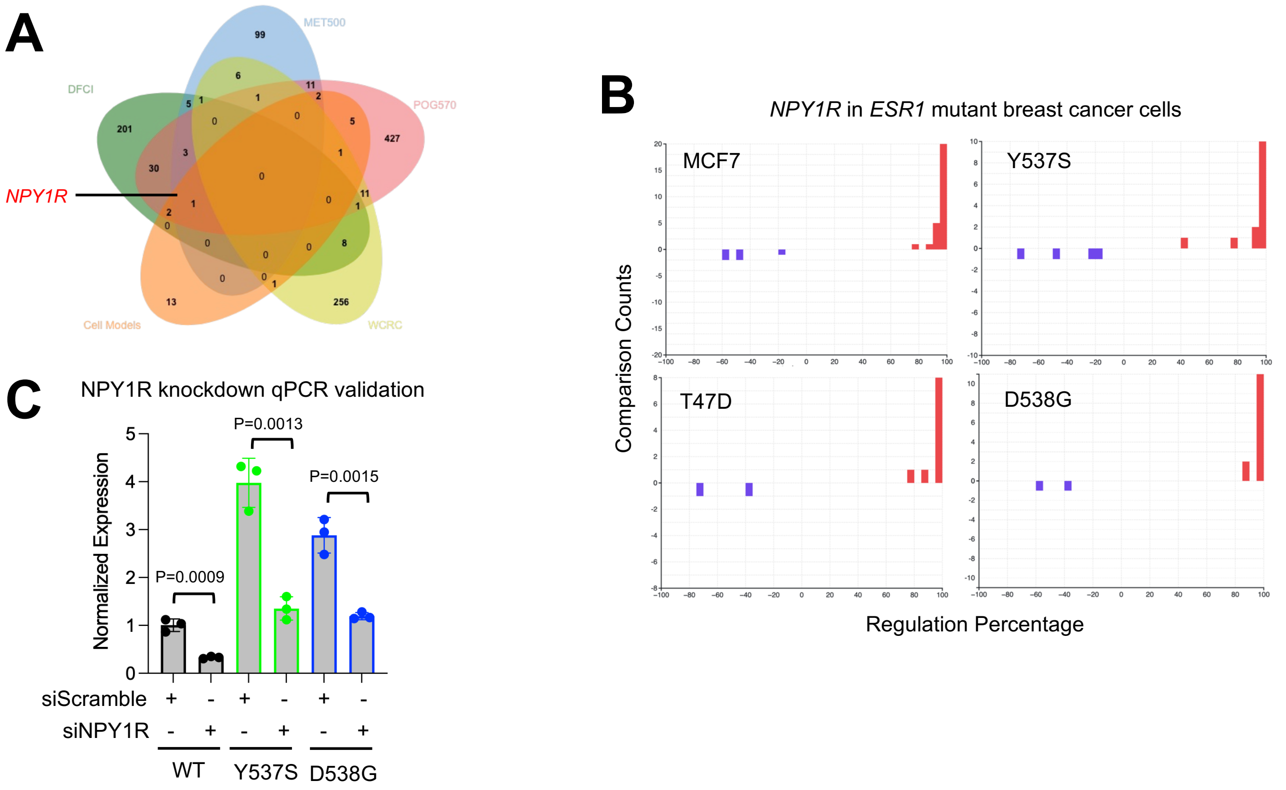


Supplementary Figure S7 (related to Fig. 6)

A. Venn diagram showing the overlap of top consistent upregulated genes in ESR1 mutant cell models cell models (from Fig. 7A) and significantly upregulated genes in ESR1 mutant tumors from four ER+ metastatic cohorts.

B. Bar plot showing the distribution of regulatory percentile of NPY1R in the selected comparisons from ESR1 mutation cell models. Percentiles are ranged between -100 to +100 and larger number indicates stronger regulation.

C. Box plots showing the relative expression of NPY1R in genome-edited MCF7 and T47D ESR1 mutant cells and WT cells in the presence of scramble and NPY1R siRNA transfection . $\Delta\Delta$Ct method was used and p values were calculated using one-way ANOVA.
